# The breast cancer pro-metastatic phenotype requires concomitant hyper-activation of ECM remodeling and dsRNA-IFN1 signaling in rare clone cells

**DOI:** 10.1101/2022.07.13.499961

**Authors:** Niccoló Roda, Andrea Cossa, Roman Hillje, Andrea Tirelli, Federica Ruscitto, Stefano Cheloni, Chiara Priami, Alberto Dalmasso, Valentina Gambino, Giada Blandano, Andrea Polazzi, Paolo Falvo, Elena Gatti, Luca Mazzarella, Lucilla Luzi, Enrica Migliaccio, Pier Giuseppe Pelicci

**Affiliations:** Department of Experimental Oncology, European Institute of Oncology, via Adamello 16, 20139 Milan, Italy; Dipartimento di Oncologia ed Emato-Oncologia, Universita’ degli Studi di Milano, via Santa Sofia 9, 20142 Milan, Italy

## Abstract

The molecular determinants of breast cancer (BC) pro-metastatic phenotype are largely unknown. Here, we leveraged lentiviral barcoding coupled to single-cell RNA sequencing to trace clonal and transcriptional evolution during BC metastatization. We showed that metastases derive from rare pro-metastatic clones that are under-represented in primary tumors. Both low clonal-fitness and high metastatic-potential are independent of clonal origin. Differential expression and classification analyses revealed that the pro-metastatic phenotype is acquired in rare cells by concomitant hyper-activation of extracellular-matrix remodeling, dsRNA-interferon signaling, and stress-response pathways. Notably, genetic silencing of single pro-metastatic genes from different pathways significantly impairs migration *in vitro* and metastatization *in vivo*, with negligible effects on cell proliferation and tumor growth. In addition, gene-expression signatures from identified pro-metastatic genes predicts metastatic progression in BC patients, independently of known prognostic factors. This study elucidates previously unknown mechanisms of BC metastatization, and provides novel prognosis predictors and therapeutic targets for metastasis prevention.

## INTRODUCTION

Breast cancer (BC) is the most commonly diagnosed cancer in women and the second cause of cancer-associated death in the female population [1], with patients’ mortality being nearly exclusively (90%) due to advanced metastatic disease [2]. The 5-year survival rate for women with metastatic BC is ∼27% and, despite the significant therapeutic progresses of the last years, metastatic BC remains mostly incurable [3].

An alternative approach to reduce BC mortality is the prevention of the advanced metastatic disease. Metastatic dissemination starts in the primary tumor (PT), where cancer cells acquire the ability to invade the surrounding tissues, migrate into the blood vessels, travel in the bloodstream, extravasate at distant sites, and ultimately form metastases [4]. In most cases, metastases appear after the surgical resection of an apparently localized disease, even after several years, as a result of the growth of clinically undetectable micro-metastases already present at diagnosis [5, 6]. Treatment of micro-metastases with combination therapy, administered in either neo-adjuvant or adjuvant setting, decreases the risk of relapse by about one third [3]. In spite of this, 30% of early-stage BC patients eventually succumb to metastatic disease [7], [8]. Of note, relapse after surgical resection is frequently preceded by the presence of circulating tumor cells [5], suggesting that metastatization is an active and long-lasting process that can be supported by micrometastases. However, a therapeutic strategy based on the specific targeting of the metastatic process is still missing, as our understanding of the molecular mechanisms of metastatization is largely incomplete.

Seminal work showed that the potential of metastasis spreading in BC significantly correlates with the degree of genetic and/or phenotypic heterogeneity in the PT [9], [10], [11]. Early studies hypothesized that specific mutations in primary breast tumors are selected by the metastatic phenotype [12]. Other studies, however, revealed that the mutational profiles of PT and matched metastases are largely overlapping, ruling out a major role of genetic mutations in the process of metastatization [13], [14].

More recent studies have suggested that metastasis spreading may be instead the consequence of the adaptation of cancer cells to unfavorable conditions, such as oxygen or nutrient deprivation and chemotherapy administration [15]. In this respect, several stress-response pathways have been implicated in the metastatic process in different experimental conditions, including the hypoxia [16], unfolded-protein [17] or antioxidant responses [18], or the dsRNA and type-I interferon signaling pathways [19]. However, how and whether activation of these pathways contributes to the acquisition of the pro-metastatic phenotype by individual cells within PT is unknow.

Lineage tracing represents an extremely powerful approach to study evolving phenotypes *in vivo*. It entails the genetic labeling of individual cancer cells with molecular barcodes and allows to trace their clonal progeny during metastatic progression [20]. Previous lineage tracing studies revealed that BC metastases are composed of a 3-to 10-fold lower number of clones as compared to their PT counterparts [21], [22], and that the least abundant clones in primary tumors dominate matched metastases across different organs [21], [22]. The molecular determinants of the disseminating pro-metastatic clone, however, have not yet been characterized.

In this study, by simultaneously tracing transcriptional states and lineages at single-cell level, we describe the transcriptional determinants of pro-metastatic cells in the growing tumors.

## RESULTS

### Pro-metastatic clones are rare and under-represented in primary tumors, and form monoclonal metastases

To identify pro-metastatic clones in growing tumors (i.e., clones in the PT that complete the metastatic cascade), we traced lineages of human BC-cells during metastatization using a library of >5 million unique barcodes (Perturb-seq guide barcode (GBC) library [23], [24]. Library complexity was estimated by the overlap of two sequencing runs (Fig.S1A, Materials and Methods). Upon viral transduction, GBCs integrate in the genome and are expressed as mRNA molecules, thus allowing parallel genomic and transcriptional lineage-tracings [23], [24].

Two independent populations of the human triple-negative breast cancer (TNBC) cell line MDA-MB-231 (reference A and B) were infected with lentiviruses expressing the GBC library and the Blue Fluorescence Protein (BFP) and, after short *in vitro* expansion (∼7 days), orthotopically transplanted into immunodeficient NSG mice (200,000 cells per mouse; 3 mice *per* each reference population; Fig. 1A). PTs were surgically removed at ∼40 days post-injection (Fig. S1B) and metastasis-infiltrated organs (lungs and liver) collected after additional ∼20 days. GBC-expressing BC cells were isolated from PTs and metastases by fluorescence-activated cell-sorting (FACS) of BFP-positive cells (Fig. S1C).

**Fig. 1.**
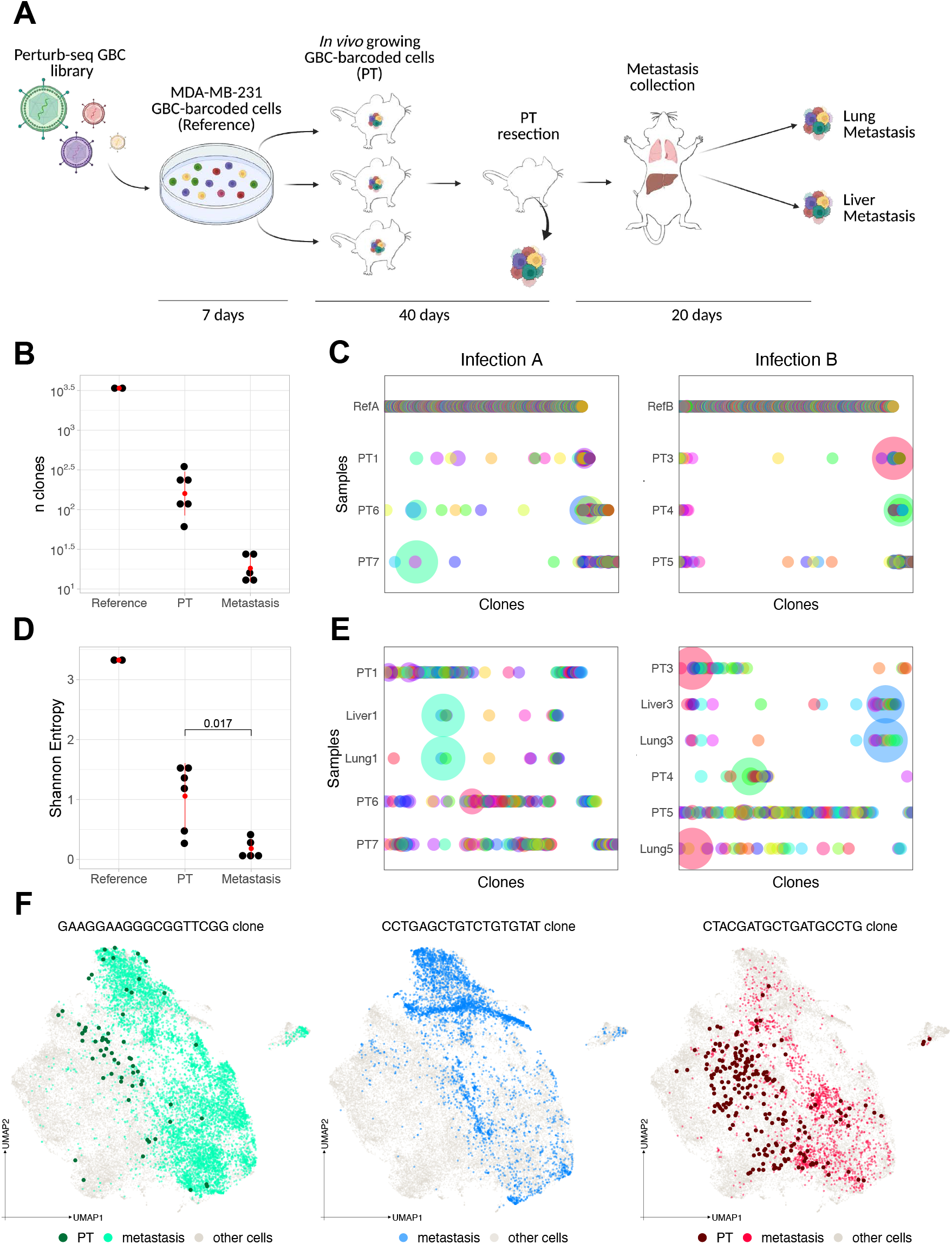
Pro-metastatic clones are rare and under-represented in primary tumors, and form monoclonal metastases. (A) Scheme of cellular GBC-barcoding of MDA-MB-231cells and *in vivo* protocol of PTs and metastasis collection. For each of two independent experiments, cells were infected *in vitro* (reference) and orthotopically transplanted in 3 NSG mice. PTs were removed at ∼40 days after injection and metastases collected ∼20 days after PT resection. GBCs were identified in tumor cells isolated from PTs and metastases by DNA-sequencing and scRNA-seq. (B) Number of clones identified in the *in vitro* reference populations, PTs and metastases. Clones were defined as the unique GBCs accounting for >90% of total reads in the corresponding sample. Standard deviation and mean are indicated in red. (C) Bubble plots of clonal frequencies in reference populations and PTs from the two independent infections. Each dot represents a clone, with radius scaled by clone frequency in the corresponding sample. Color codes are related between reference and PT populations and between different mice. (D) Clonal distribution Shannon Entropy (SE) of reference populations, PTs and metastases. SE was calculated by considering the frequency of all clones in the corresponding samples. Blue bars indicate standard deviation; *in vitro* reference n = 2; PTs n = 6; metastases n = 5; Wilcoxon-Mann-Whitney test (PTs vs metastases) p = 0.017. Standard deviation and mean are indicated in red. (E) Bubble plots of clonal frequencies in PTs and matched metastases from the two independent infections (as in C). Color codes are related between PT and metastasis and between different mice. (F) UMAP representation of scRNAseq data of all 6 PTs and 5 metastases showing the distribution of cells of the dominant pro-metastatic clones. Dominant pro-metastatic clone cells in PT and metastatses are indicated with different colors. PT cells are represented with a darker color and bigger size to facilitate their identification. The GBC identity of each pro-metastatic clone is shown above. In grey, we reported all the other cells.

To infer clonal dynamics during PT growth, we sequenced genome-integrated GBCs from the 2 *in vitro* references and the 6 matched PTs. Clones were defined as the unique GBCs accounting for >90% of total reads. Numbers of clones were dramatically reduced in PTs, as compared to *in vitro* references: ∼170 clones (C.I.95 = 170 ± 65) in each PT and ∼3,400 clones (C.I.95 = 3,380 ± 56) in each reference (Fig.1B-C). Reduced clonal complexity was confirmed by the decrease of clonal-distribution Shannon Entropy (SE) from *in vitro* references to PTs (Fig. 1D). Highly comparable clonal-frequencies were inferred from single-cell RNA sequencing (scRNA-seq) in 4 out of 6 PTs (Fig. S1D).

Clonal dynamics of metastatization were then investigated by analysis of scRNA-seq from 5 metastases: lung and liver metastases matched to PT1 or PT3 (Dataset_1 or _3), and lung metastasis matched to PT5 (Dataset_5). Clonal complexity was even lower in metastases than in PTs, with ∼10 clones per metastasis (C.I.95 = 11 ± 4, Fig. 1B and 1E) and significantly lower clonal-distribution SE (SE = 1.06 and 0.18 in PTs and metastases, respectively; p=0.0093; Fig. 1D), suggesting that only a small fraction of the PT clones can colonize distant sites. Strikingly, a single clone accounted for most cells in each metastasis (C.I.95 = 92±7% of total cells; dominant pro-metastatic clones), with the other clones showing different size distribution (C.I.95 = 0.44 ± 0.01% cells per clone; minor pro-metastatic clones) (Fig. 1E). In line with previous studies [21], [22], the dominant pro-metastatic clones were highly under-represented in the corresponding PTs, accounting for 1.65·10^−3^, 1.4·10^−7^ and 3.5·10^−2^ of all PT1, PT3 and PT5 cells, respectively). In addition, matched lung and liver metastases were populated by the same pro-metastatic clone (Fig. 1F).

Thus, metastatic progression in BC is mainly driven by rare pro-metastatic clones, which are under-represented in the corresponding PTs (i.e., display low fitness in the PT microenvironment) and colonize different organs.

### Low clonal-fitness and high metastatic potential are lineage-independent properties of the pro-metastatic clones

We next investigated whether low clonal-fitness and high metastatic potential are intrinsic properties of pro-metastatic clones or are instead acquired during tumor growth. To this end, we took advantage of rare clones (96 out of the 990 identified across all PTs) that expanded in multiple PTs from the same infection (common clones): 56 across PT1, 6 and 7; 50 across PT3, 4 and 5.

Clonal frequencies in reference populations spanned a narrow range (Fig. 2A), suggesting homogeneous fitness *in vitro*. Conversely, *in vivo* clonal frequencies spanned a broad range (Fig. 2B), with <10 clones accounting for >75% of the PT biomass (C.I.95 = 9.00 ± 5.49; see also Fig. 1B-D), suggesting that clones acquire heterogenous fitness during tumor growth. Strikingly, common clones showed different (Fig. 2C) and minimally-correlated (Fig. 2D) frequencies across most PTs, suggesting that clonal fitness is acquired during tumor growth. Analysis of pro-metastatic potential of common clones revealed the presence of 11 clones common to PT3 and PT5 that disseminated only in either PT3-(5 clones) or PT5-(6 clones) metastases. Thus, the same clone can be either pro- or non-metastatic according to the specific PT-contexts, suggesting that the metastatic potential is also acquired during tumor growth. A peculiar case was represented by the clone CTACGATGCTGATGCCTG. This clone contributed to 3.5% and 87% cells in PT5 and PT3, respectively (Fig. 2E), in line with the clonal fitness lineage-independency. However, this clone turned out to be dominant pro-metastatic in the dataset 5 and minor pro-metastatic in the dataset 3 (Fig. 2E), confirming that the metastatic potential of a clone can be extremely different in distinct PT-contexts.

**Fig. 2.**
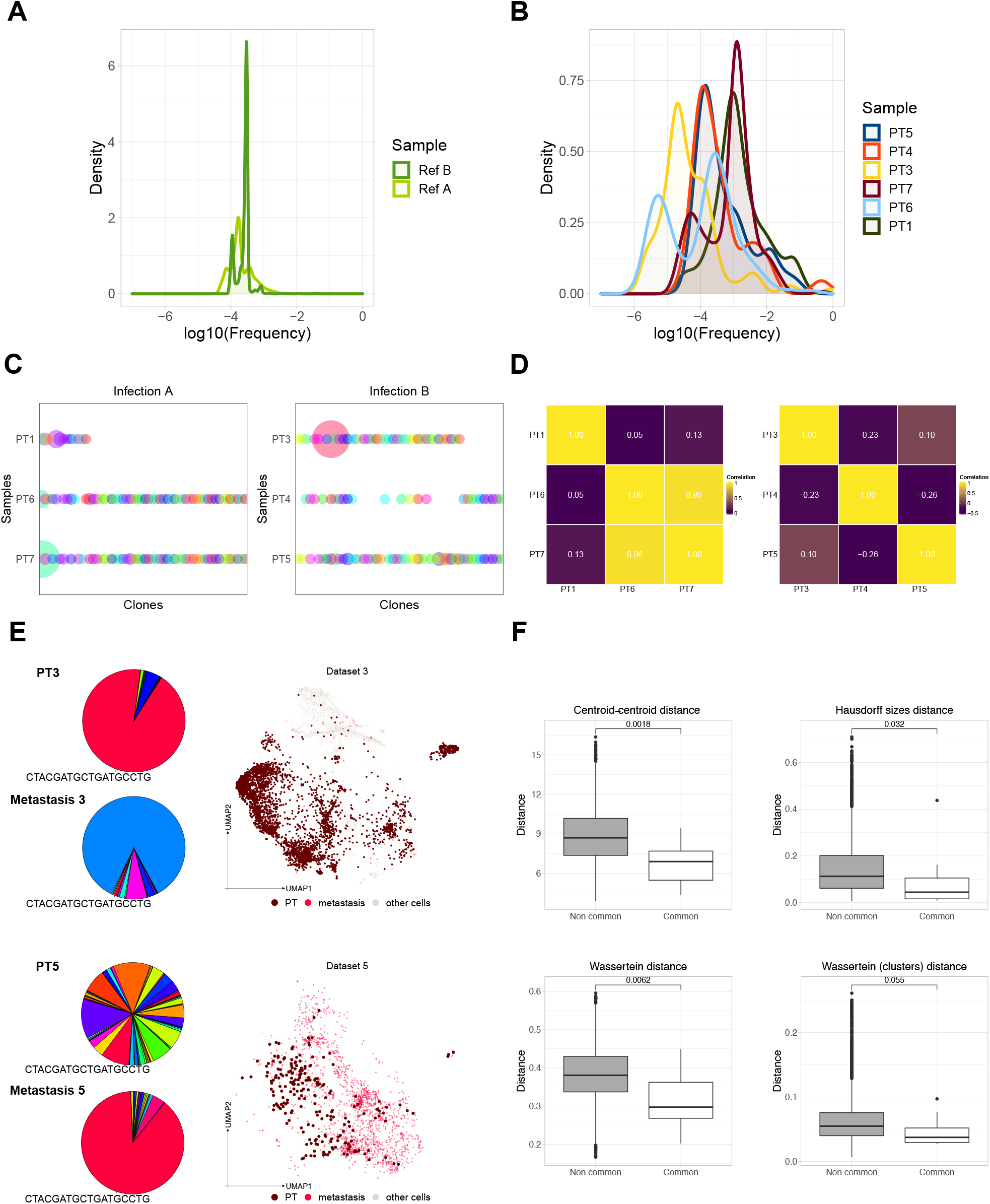
Low clonal-fitness and metastatic potential are lineage-independent properties of the pro-metastatic clones. (A,B) Log(10)-frequency distribution of clones in the two *in vitro* references (A) and six PTs (B), as indicated. Density defines per each value of clonal frequency the inferred number of clones with that frequency. (C) Bubble plots of common-clone frequencies in reference and PT cell-populations from the two independent infections (as in Fig. 1C and 1E). (D) Pairwise Pearson correlation of common-clone frequencies across multiple PTs in the two infections. (E) UMAP representation showing the CTACGATGCTGATGCCTG clone in Dataset5 and Dataset3. The CTACGATGCTGATGCCTG cells in the PT are represented with a darker color and bigger size. CTACGATGCTGATGCCTG cells in metastases are colored with the specific clone-color. In grey, we reported all the other cells of the corresponding PT (other cells). (F) Pairwise transcriptional distances (see Materials and Methods) among clonal populations belonging to either common or non-common clones. Wilcoxon-Mann-Whitney test p-values are reported.

We next investigated whether the transcriptional state of a cell is influenced by its lineage. To this end, we computed pairwise transcriptional-distances among all clones across PTs (only clones with ≥ 20 cells in their respective PT were considered; C.I.95 = 18.33 ± 9.30 clones per PT) and compared distances between common (i.e., lineage-related) and unique (i.e., non-lineage-related) clones. We chose metrics able to quantify distance among either probability distributions (Wassertein distance) or metric spaces (Hausdorff distance, centroid-centroid distance). Independently of the chosen metrics, common clones in different PTs were on average more similar than unique clones (Fig. 2F, Materials and Methods), suggesting that the transcriptional state of a given cell is significantly influenced by its clonal derivation.

Together, these data suggest that the overall transcriptional state of BC cells is an intrinsic property of each clone. Nevertheless, diverging functional phenotypes can be acquired during tumor growth *in vivo*, including clonal fitness and metastatic potential.

### Dominant pro-metastatic clones are characterized by the same intra-clonal heterogeneity of all other PT clones

Intra-tumor heterogeneity is considered a major driver of metastatization, as it provides the cellular background for the selection of specific pro-metastatic phenotypes [25]. Thus, we investigated whether dominant pro-metastatic clones are enriched in specific regions of the reduced gene-expression space (i.e., transcriptional clusters).

The 37,801 cells of our dataset (22,313 from PTs and 15,488 from matched metastases; Fig. 3A) were grouped in 11 transcriptional clusters (Fig. 3B). Cell-cycle phase-scoring assigned ∼51% of PT cells to either S or G2/M and ∼49% to G0/G1 (Fig. 3C). Consistently, immunochemistry on PT paraffin-embedded sections revealed ∼47% positivity for the Ki67+ proliferation marker (Fig. S2A). Differences in cell cycle distribution segregated markedly across clusters (Fig. 3C), with clusters 1, 5, 8 and 9 showing >75% S-G2/M cells (C.I.95 = 88.8 ± 11.2%; highly-proliferative Group 1 clusters), and clusters 2, 3, 4, 6, 7, 10 and 11 <65% S-G2/M cells (C.I.95 = 33.8 ± 18.1%; poorly-proliferative Group 2 clusters) (Fig. S2B).

**Fig. 3.**
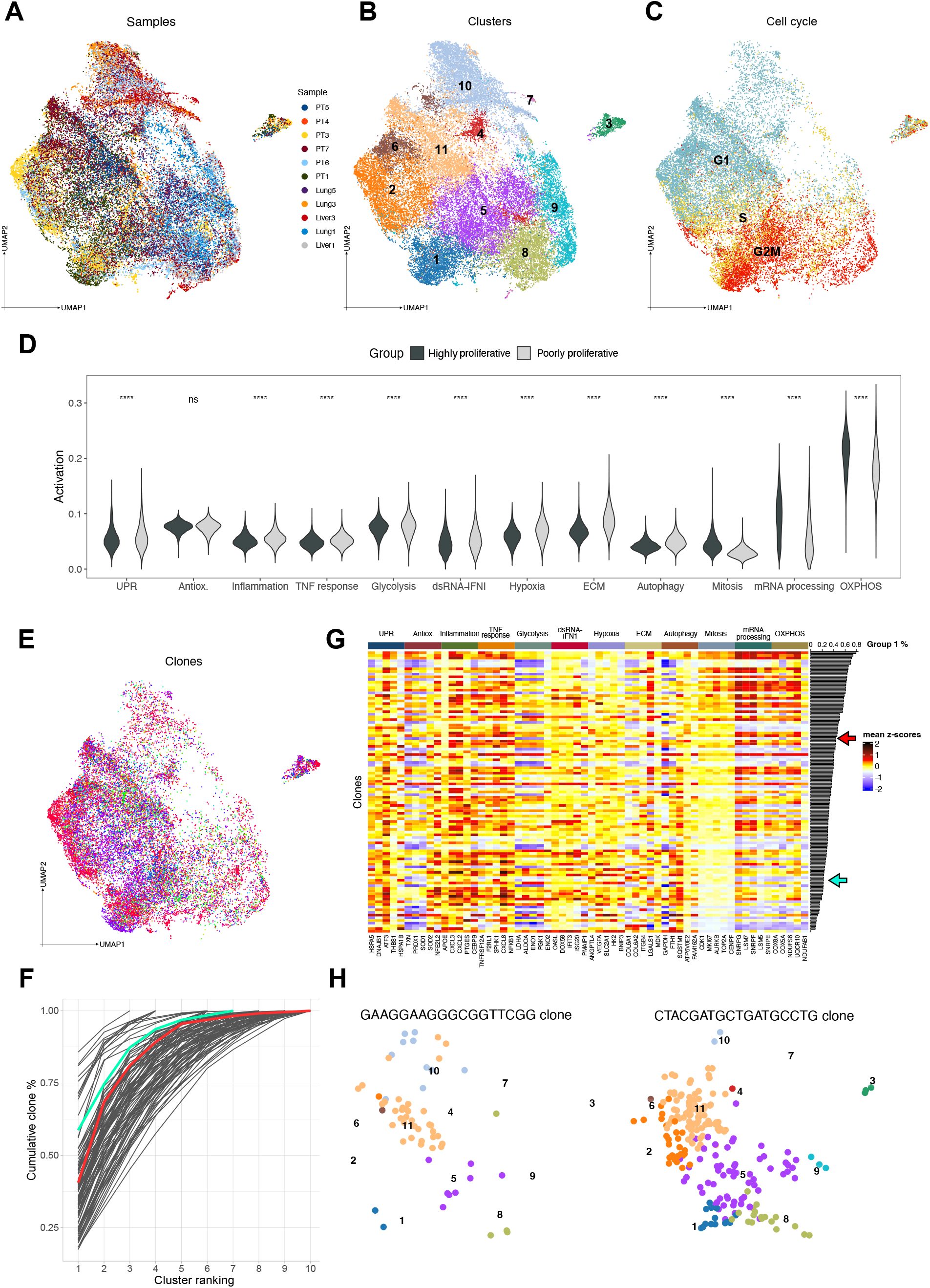
Dominant pro-metastatic clones are characterized by the same intra-clonal heterogeneity of all other PT clones. (A) UMAP representation of 37,801 cells colored by sample. (B) UMAP representation of 37,801 cells colored by Louvain clustering. (C) UMAP representation of 22,313 single cells from the 6 PTs colored by cell cycle phase assignment. (D) Violin plots showing expression of gene signatures associated to mitosis (GOBP_MITOTIC_NUCLEAR_DIVISION), mRNA processing (GOCC_SPLICEOSOMAL_TRI_SNRNP_ COMPLEX), OXPHOS (GOBP_OXIDATIVE_PHOSPHORYLATION), UPR (GOBP_CELLULAR_RESPONSE_TO_UNFOLDED_ PROTEIN), hypoxia (GOBP_CELLULAR_RESPONSE_TO_ OXYGEN_LEVELS), autophagy (GOCC_ AUTOPHAGOSOME), dsRNA-IFN1 signaling (GOBP_CELLULAR_RESPONSE_TO_DSRNA), TNF-α response (GOBP_RESPONSE_TO_TUMOR_NECROSIS_FACTOR), ECM remodeling/ interaction (GOCC_COLLAGEN_ CONTAINING_EXTRACELLULAR_MATRIX), glycolysis (HALLMARK_GLYCOLYSIS), oxidative-stress response (GOBP_RESPONSE_TO_OXIDATIVE_STRESS), and inflammation (GOBP_RESPONSE_TO_MOLECULE_OF_ BACTERIAL_ORIGIN) in all Group 1 (1, 5, 8, and 9) *vs* all Group 2 (2, 3, 4, 6, 7, 10, and 11) clusters (Wilcoxon-Mann-Whitney test; **** p < 0.0001). (E) UMAP representation of 22,313 cells from the 6 PTs colored by clone. Clones are indicated by different colors. (F) Cumulative clone percentages (each clone is represented as a line). The cumulative percentage of a given clone (y-axis) was calculated by progressively adding the cluster frequencies of that clone, sorted in ascending order (x-axis). PT1 and PT5 dominant pro-metastatic clones are colored in green and red, respectively. (G) Heatmaps interrogating gene expression across clones of 5 representative genes (i.e., either assigned by over-representation analysis to the corresponding pathway or manually curated (e.g., VEGFA and HK2)) per each of the 12 pathways identified by cluster analyses in the PT clones. Clones are ranked from top to bottom according to decreasing fraction in Group 1 clusters (Group 1 %). PT1 and PT5 dominant pro-metastatic clones are indicated by the green and red arrows respectively. (H) UMAP representation of the dominant pro-metastatic clones from PT1 (left panel) and PT5 (right panel). Cells are colored by Louvain clustering.

To analyze transcriptional features of single clusters, we performed marker-gene analysis (FDR<0.1; **Supplementary Table 1** and Fig. S2C) and used the top 50 marker-genes *per* cluster (total of 1,715 non-redundant genes) to identify over-represented gene-sets in different collections of the Molecular Signatures database (MSigDB; gene ontology (GO) and curated collections; results of all analyses are provided in **Suppl. Table 1**). The top 1-2 over-represented gene-sets per cluster clearly identified putative functions of Group 1 and 2 clusters. Group 1 Clusters showed enrichment of genes involved in mitotic progression (e.g., CDK1, AURKB, MKI67, TOP2A, CENPF/W; Clusters 1, 5 and 8), oxidative phosphorylation (OXPHOS; e.g., COX8A/7A2/5A, UQCRH/10; Clusters 5, 8 and 9) and mRNA processing (e.g., SNRPG/F/E/D1/D3, LSM3/5/7; Clusters 5, 8 and 9). Group 2 clusters, instead, showed enrichment of genes involved in pathways previously implicated in the regulation of metastasis: i) integrated cellular responses to stress: unfolded protein response (UPR; e.g., HSPA5 (BiP), DNAJB1 (HSP40), ATF3, THBS1, and HSPA1B (HSP70.2)), autophagy (e.g., FTH1,

FTL, SQSTM1), hypoxia (e.g., NRDG1, BNIP3, HK2), and tumor necrosis factor-α (TNF-α; e.g., TNFRSF12A, SPHK1, F2RL1) or antioxidant responses (NFE2L2, SOD1, TXN, and PRDX1) (Clusters 2, 3, 6 and 7); ii) dsRNA (e.g., OASL, DDX58 (Rig-1), IFIH1 (MDA5)) and IFN1 (e.g., ITIF1/2/3, ISG15/20) signaling (Cluster 4); iii) inflammation (e.g., CEBPB, CXCL2/3/8, IL7R; Cluster 11); iv) ECM remodeling/interaction (e.g., ADAMSTS1, FN1, S100a11/6; Clusters 10 and 11). As well, glycolysis was also enriched in Cluster 6; glycolysis and OXPHOS in Clusters 3 and 7. Similar patterns of pathway over-representation were obtained upon cell-cycle regression of the scRNA-seq dataset (**Suppl. Table 2**).

We then investigated differential expression of curated gene-signatures for each of the 12 over-represented pathways. Each signature was scored across single clusters (Fig. S2D) or between all Group 1 vs all Group 2 clusters (Fig. 3D). Strikingly, cell proliferation, mRNA processing and OXPHOS gene-signatures were significantly over-expressed in Group 1 clusters, while UPR, hypoxia, glycolysis, autophagy, TNF-α response, dsRNA-IFN1 signaling, inflammation, and ECM remodeling/interaction were over-expressed in Group 2 Clusters (Fig. S2D and Fig. 3D).

As orthogonal approach, we clustered genes with co-varying expression-patterns in the reduced gene expression space (Hotspot method; Materials and Methods) and identified 7 gene-modules (GMs; Fig. S2E and **Suppl. Table 3**), which showed enrichment of the same pathways as found with cluster markers (**Suppl. Table 1**). Strikingly, GMs associated to mitotic cell-division and mRNA processing were over-expressed in Group 1 clusters, while GMs associated to stress response pathways (UPR and hypoxia), dsRNA-IFN1 signaling, glycolysis, and ECM remodeling/interaction were over-expressed in Group 2 clusters (Fig. S2F). Thus, clusters and GMs showed increased expression of one of two mutually exclusive transcriptional programs: highly- or poorly-proliferative, the latter endowed with variable activation of different stress response pathways, ECM remodeling/interaction, and glycolysis.

We next investigated the relationships between PT clusters and clones (only clones with ≥ 20 cells in their respective PT were considered; Fig. 3E). Each cluster was populated by multiple clones (C.I.95 = 85.30 ± 10.10) and each clone was composed of multiple clusters (C.I.95 = 8.12 ± 0.32, Fig. S3A). Intra-clonal heterogeneity was quantified using cluster distribution SE, a measure that in our dataset can take all values between 0 (when a clone populates a single cluster: SE=0) and 1.04 (when a clone is distributed uniformly across all 11 clusters: SE=log1011). Cluster distribution SE displayed an average value of ∼0.7 (C.I.95 = 0.72 ± 0.03) and a distribution skewed towards higher values, suggesting a high degree of intra-clonal heterogeneity (Fig. S3B). On average, however, a reduced number of clusters (∼3; C.I.95 = 3.22 ± 0.33) accounted for >75% of cells in each clone (Fig 3F and S3C). Specifically, ∼80% of clones were mainly enriched in Group 2 clusters (i.e., contained >50% of their cells in Group 2 clusters), while remaining clones were mainly enriched in Group 1 clusters (Fig.3G). Concordantly, poorly-proliferative Group 2 clones displayed higher expression of genes related to cellular response to stress, autophagy, and dsRNA-IFN1 signaling, while highly-proliferative Group 1 clones displayed higher expression of genes related to mitotic cell division, mRNA processing and OXPHOS (Fig. 3G). Thus, each clone retains most of the potential of the entire PT to generate transcriptional heterogeneity, yet it is canalized towards specific transcriptional states.

Finally, we analyzed cluster distribution of PT1 and PT5 dominant pro-metastatic clones (63 and 222 cells, respectively). Both clones contained multiple clusters (7 and 10, respectively) (Fig. 3H), and showed a cluster-distribution SE within or immediately below the SE range of the other clones (0,68 and 0,56 for PT1 and PT5 clone, respectively) (Fig. S3B). As observed for all other PT clones, 3 clusters accounted for >75% of all cells in both dominant pro-metastatic clones (i.e., clusters 11, 10, and 5 in PT1 clone; clusters 11, 5, and 2 in PT5 clone; Fig. 3F). Dominant pro-metastatic cells were enriched for Group 2 clusters (78 and 56% cells in PT1 and PT5 clone, respectively) and G0-G1 phase (65 and 61% cells, respectively; Fig. S3D), similarly to ∼80% non-metastatic clones. Thus, PT1 and PT5 dominant pro-metastatic clones are enriched for poorly-proliferative cellular states and are as transcriptionally heterogeneous as the other clones. Together, these data argue against the hypothesis that the metastatization process selects clones with homogeneous transcriptional states.

### Dominant pro-metastatic clones are distinguished by up-regulation of genes that predict disease progression in BC patients

To investigate distinguishing transcriptional-features of dominant pro-metastatic clones, we first performed Differential Expression (DE) analyses (**Suppl. Table 4**) in four different settings: i) pro-metastatic *vs* all the other cells in separate *(‘Single-PT’* comparisons) or pooled *(‘Pooled-PTs’*) PT1 and PT5; ii) pro-metastatic cells *vs* their ‘nearest’ cells of separate (*‘Single-NN’*) or pooled (*‘pooled-NN’*) PT1 and PT5; iii) pro-metastatic *vs* the other cells of the two clusters (5 and 11) most enriched in pro-metastatic cells (*‘Pooled-cluster’*); and iv) dominant (PT5) *vs* minor (PT3) pro-metastatic cells from the common clone CTACGATGCTGATGCCTG (*‘Common-clone’*). These comparisons yielded 9 DEGs lists (**Suppl. Table 4**), with variable degree of overlap (Jaccard Index, J.I., Fig. 4A). In particular, *‘Single-PT’* and S*ingle-NN*’ comparisons identified a large set of overlapping DEGs for each pro-metastatic clone (J.I. =0.30 and 0.48 for PT1 and PT5, respectively; Fig. 4A and S4A), while DEGs from the two pro-metastatic clones showed minimal overlap, suggesting the existence of clone-specific pro-metastatic phenotypes (J.I. = 0.02 and 0.01 between PT1 and PT5 ‘*single-PT*’ and ‘*single-NN*’ DEGs, respectively; Fig.4A and S4B). PT1- and PT5-clone DEGs are shown in Fig. 4B (‘*single-PT*’) and Fig. S4C (‘*single-NN*’). Overlaps among the other DEG lists were variable, but overall limited (J.I. ≤0.26; Fig.4A), suggesting that different comparisons captured different components of the pro-metastatic phenotype. To confirm these results with an independent approach, we exploited machine learning models to classify pro-metastatic cells vs all the other cells from their transcriptional features (logistic regression and extreme gradient boosting models, fed with either Hyper-Variable Genes (HVGs), PCA space or scVI cell embeddings). Predictions were evaluated with training and test sets derived within separate or pooled PT1 and PT5, and across PTs (i.e., one PT as training and the other as test set; Fig. S4D). As expected from DE analysis, classification performance was consistently higher for separate PT1 and PT5 predictions and dropped progressively in pooled-PTs and across-PTs settings (Fig. S4E). Within each PT, the best model correctly identified >75% pro-metastatic and >95% other cells (F1-score = 0.74 and 0.58 for PT5 and PT1, respectively, Fig. 4C). Notably, for each PT clone, genes prioritized with this classification approach (**Suppl. Table 5**; Materials and Methods) significantly overlapped with DEGs identified in the *‘single-PT’* and ‘*single-NN*’ settings (Fig. 4D). Thus, independent methods demonstrated the existence of distinguishing transcriptional features of pro-metastatic cells, which however differ across the two pro-metastatic clones.

**Fig. 4.**
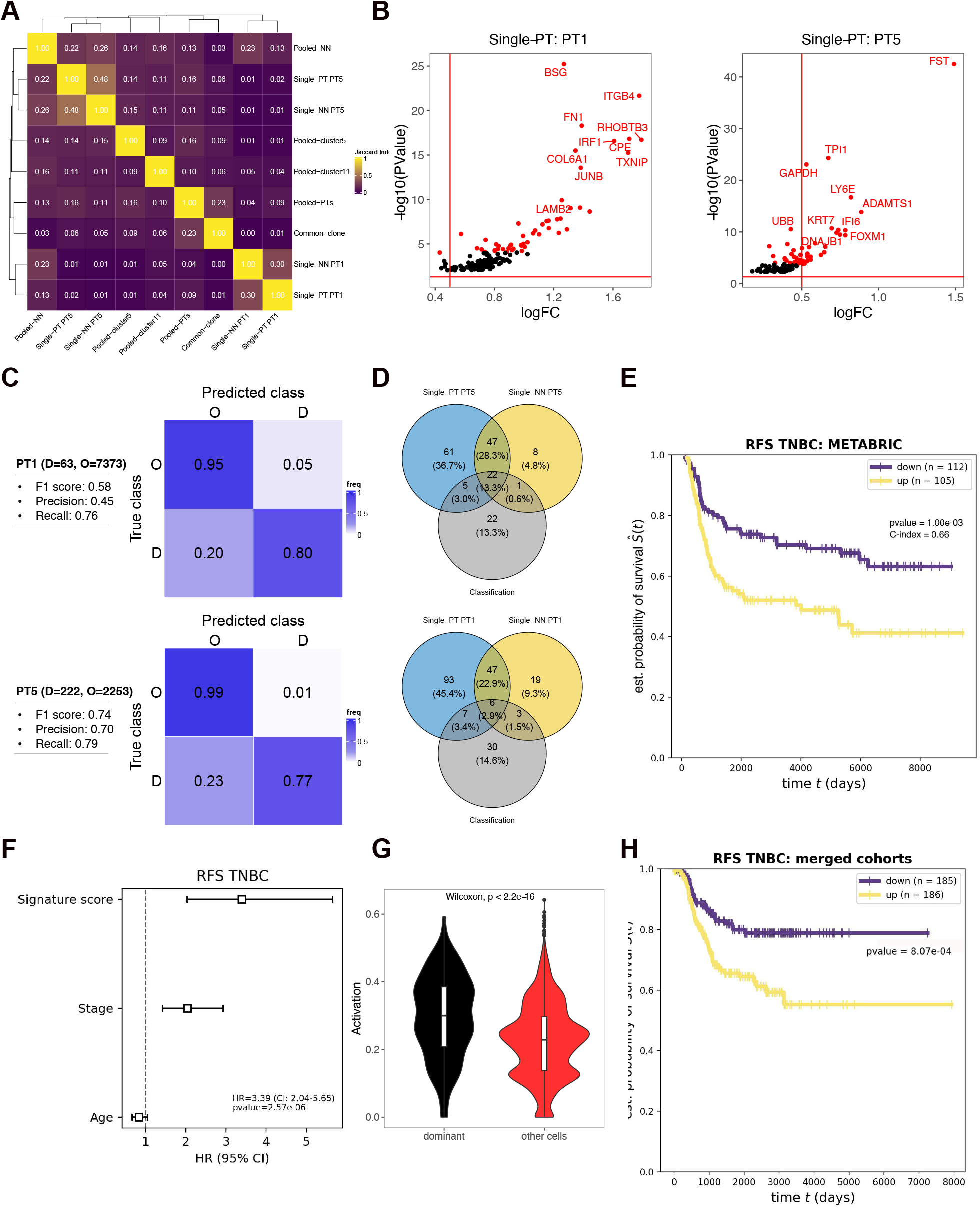
Dominant pro-metastatic clones are distinguished by up-regulation of genes that predict disease progression in BC patients. (A) Pairwise similarity matrix showing the degree of overlap (quantified by Jaccard Index) among all DE lists. Both rows and columns were hierarchically clustered. (B) Volcano plot representing dominant pro-metastatic clones’ DEGs identified in the ‘single-PT’ setting (left PT1 and right PT5, respectively). Per each DEG, we reported the corresponding logFC (x-axis) and - log10(Pvalue) (y-axis). The top 50 DEGs (ranked by FDR) are colored in red. (C) Classification performance of the two best models from the “separate PT” scenario: logistic regression fed with PCA space (PT1, upper panels); and logistic regression fed with HVGs (PT5, lower panels). For each PT we report the cell counts of the two considered classes: D (dominant pro-metastatic cells) and O (other cells). For each model, we report precision (the fraction of correct D calls over the total D calls), recall (the fraction of correct D calls over the total D cells), and F1 score (the harmonic mean between precision and recall) evaluation metrics. Each model performance is visualized with a confusion matrix, where the “True” class represents the class to which a cell actually belongs and the “Predicted class” represents the class to which each PT cell has been assigned by the classifier. Each confusion matrix is row-normalized. (D) Venn diagram representing the top 50 genes in the ‘single-PT’ and ‘single-NN’ DE settings together with the top 50 genes identified through the classification approach. (E) TNBC RFS signature performance in the METABRIC cohort. The figure shows both the Kaplan-Meier curves (with the related log-rank test p-value) and the CI of the best survival model employing the TNBC RFS signature genes as predictors (i.e., the multivariate CPH model, see Materials and Methods). The y-axis reports the estimated probability of survival (Ŝ) over time (i.e., numbers of subjects surviving divided by numbers of patients at risk), while the x-axis reports survival time (in days). (F) Multivariate analysis of the TNBC RFS gene signature score, together with other clinical factors. (G) Violin plots showing the expression of the RFS-TNBC signature in dominant pro-metastatic (dominant) vs other cells from pooled PT1 and PT5 (Wilcoxon-Mann-Whitney test, p-value is reported). (H) TNBC RFS signature performance in the validation dataset (multiple merged cohorts, see Materials and Methods). The figure shows the Kaplan-Meier curves with the related log-rank test p-value.

To obtain a preliminary validation of the output of these analyses, we investigated the capacity of the identified pro-metastatic genes to predict BC-patient prognosis. We selected the top 50 genes from each DE and classification approach (301 unique genes) and performed survival analysis on a cohort of 1,394 BC patient samples with complete expression- and clinical-data annotations at diagnosis (METABRIC). Overall Survival (OS) and Relapse-Free Survival (RFS) were considered separately for TNBC patients and for patients from all BC subtypes. For all four survival tasks (OS and RFS in TNBC or all patients), we observed a significant enrichment of clinically-relevant genes within our selected gene pool (Hazard Ratio (HR)>1 and Concordance index (CI)>0.5, from univariate Cox Proportional Hazard (CPH) regression), as compared to random (Fig. S4F; Materials and Methods). We then explored the extent of statistical interactions among candidate pro-metastatic genes, searching for prognostic gene signatures. Briefly, for each survival task we performed multivariate survival analyses with CPH regression and gradient-boosted models combined with different feature-selection procedures (univariate CPH regression, forward selection and elastic nets). The best performing gene-signature for each task successfully predicted patient survival with improved performance over single genes, and independently of other well-characterized risk factors (stage, age, and subtype; Fig.4F and S5C, **Suppl. Table 6**).

The 4 signatures (26- and 5-genes for OS on all BC and TNBC patients, respectively; 11- and 7-genes for RFS on all BC or TNBC patients, respectively; Fig. S5A) included 45 unique genes, with only 4 genes present in >1 signature. Most notably, each signature included genes obtained from PT1 and PT5 lists, suggesting that the distinct transcriptional features of the two pro-metastatic clones represents distinct aspects of the same pro-metastatic phenotype, including, for example, space in the tumor tissue or time during tumor progression. The presence of different transcriptional programs across distinct pro-metastatic clones was previously reported in a model of pancreatic ductal adenocarcinoma [26].

Consistently with our experimental system, the 7-gene signature predicting RFS in TNBC patients showed the best prognostication performance (CI= 0.66; HR= 3.39) and was significantly upregulated in pro-metastatic cells, as compared to their non-metastatic counterparts (Fig. 4H). We validated the prognostic value of the two best-performing signatures (RFS in TNBC patients and OS in all BC patients, respectively) in multiple independent patient cohorts (Materials and Methods). While neither of the two significantly stratified patients in separated cohorts (Fig. S5D-E), both succeeded upon cohorts merging (Fig. 4I and S5F).

These data demonstrate that pro-metastatic cells are distinguished by increased expression of a pool of specific and clinically-relevant genes, which can be used to derive molecular predictors of BC prognosis.

### Rare cells of the pro-metastatic clones concomitantly hyper-activate ECM remodeling/interaction, dsRNA-IFN1 signaling and stress responses (hypoxia, unfolded protein, and antioxidant)

We next investigated cellular functions and signaling pathways associated with the identified putative pro-metastatic genes. We performed pathway over-representation analyses using the top 50 genes of each dominant pro-metastatic clone obtained from either DE (‘*Single-PT*’ setting) or classification approaches in separate PTs (**Suppl. Tables 7**,**8**). Most notably, while the two methods yielded only-partially overlapping top-ranked genes (Fig. 4D), they returned similar over-represented pathways. The top 1-2 gene-sets of each clone identified ECM remodeling/interaction (divided in ECM deposition (FN1, COL6A1, and COL6A2), adhesion (ITGB4 and ITGA10) and degradation (ANGPTL4, BSG, and CTSD; Fig.5A)) and dsRNA-IFN1 signaling (Fig.5B) as distinguishing pathways of PT1 and PT5 pro-metastatic clones, respectively. Of note, investigation of lower-ranked pathways showed enrichment of ECM remodeling/interaction and dsRNA/IFN1 signaling in the other (PT5 or PT1) pro-metastatic clone, hypoxia response in both clones, antioxidant response in PT1 and UPR in PT5 pro-metastatic clones respectively (Fig.5A-B and **Suppl. Tables 7**,**8**). Strikingly, manually curated gene-signatures of ECM remodeling/interaction, dsRNA-IFN1 sgnaling, and hypoxia were over-expressed in both PT1 and PT5 pro-metastatic clones, as compared to Group 1 and, in several cases, Group 2 clones (Fig. 5C-D). Ultimately, gene-signatures related to UPR was significantly over-expressed PT5 pro-metastatic cells, while we did not find upregulation of antioxidant response in the PT1 pro-metastatic clone, consistent with the presence of a smaller subset of upregulated antioxidant-related genes in this clone (Fig. 5C-D).

**Fig. 5.**
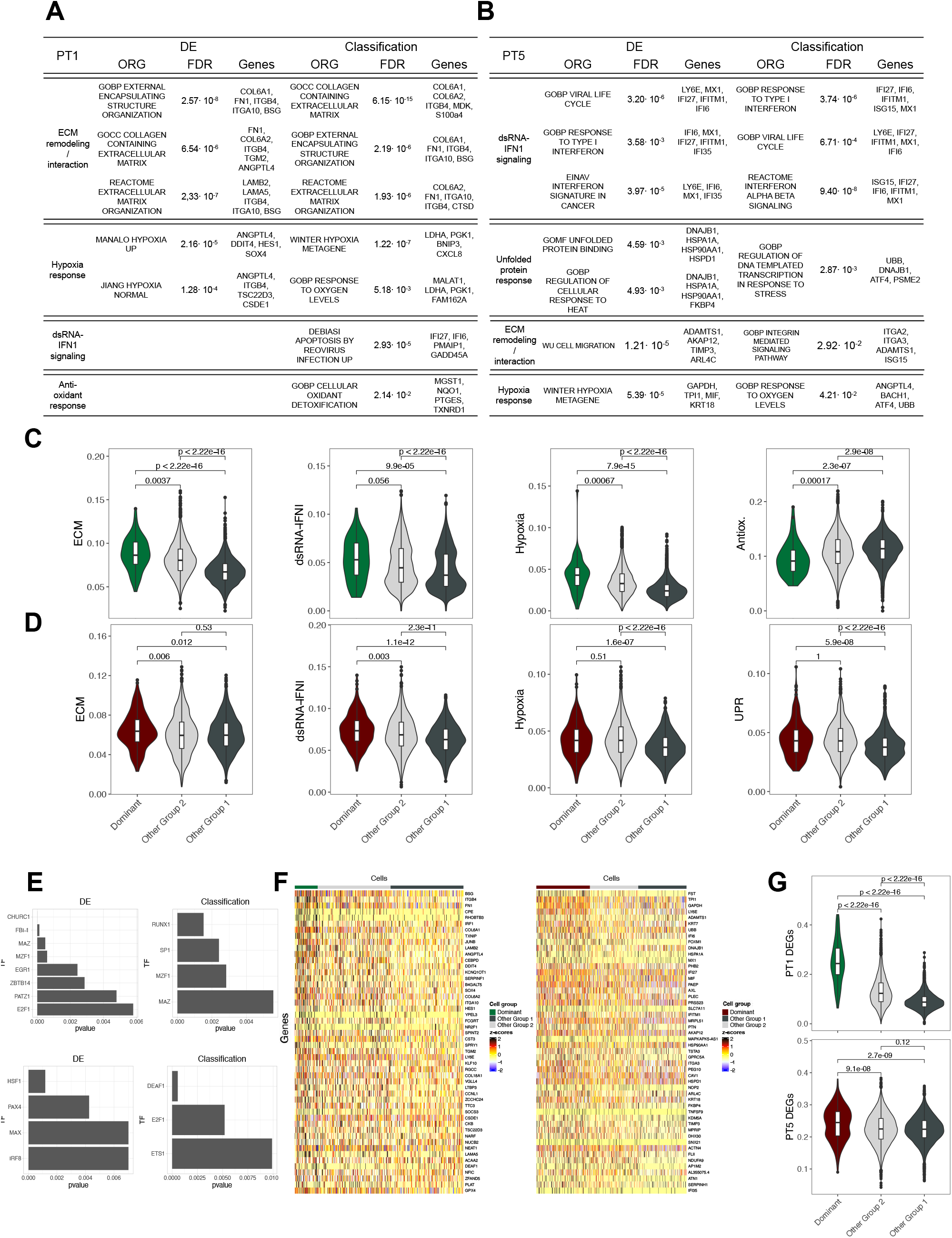
Rare cells of the pro-metastatic clones concomitantly hyper-activate ECM remodeling/ interaction, dsRNA-IFN1 signaling and stress responses (hypoxia, unfolded protein, and antioxidant) **(A-B)** Schematic representation of the functional pathways enriched in PT1 (A) and PT5 pro-metastatic clone (B). Per each clone, we report the identified functional pathway, the over-represented gene-sets (ORGs) identified by either DE or classification, the FDR associated to each ORG, and 4-5 representative genes per ORG. **(C)** Violin plot showing the expression of previously established gene signatures associated to ECM remodeling/interaction (GOCC_COLLAGEN_CONTAINING_EXTRACELLULAR_MATRIX), dsRNA-IFN1 signaling (GOBP_RESPONSE_TO_TYPE_I_INTERFERON), hypoxia response (MANALO_HYPOXIA), and antioxidant response (GOBP_CELLULAR_OXIDANT_DETOXIFICATION) in dominant pro-metastatic (Dominant) vs all the other cells (Other) in PT1 divided in Group 1 and 2. **(D)** Violin plot showing the expression of previously established gene signatures associated to ECM remodeling/interaction (GOMF_INTEGRIN_BINDING), dsRNA-IFN1 signaling (GOBP_RESPONSE_TO_TYPE_I_ INTERFERON), hypoxia response (GOBP_CELLULAR_RESPONSE_TO_HYPOXIA), and UPR (GOBP_CELLULAR_ RESPONSE_TO_TOPOLOGICALLY_INCORRECT_PROTEINS) in dominant pro-metastatic (Dominant) vs all the other cells (Other) in PT5 divided in Group 1 and 2. **(E)** Bar plot of the TFs identified by the PASTAA algorithm from the DE (left panels) or classification (right panels) genes in PT1 (upper panels) and PT5 (lower panels) pro-metastatic clones. **(F)** Heatmap showing the expression of the top 50 DEGs (identified in the ‘single-PT’ setting) in single cells of the PT1 and PT5 dominant pro-metastatic clones and in 400 randomly-sampled non-metastatic cells from the same PT (200 cells were sampled from Group 1 non-metastatic clones and 200 from Group 2 non-metastatic clones). Each column corresponds to a cell and each row to a gene. Genes are ranked by increasing FDR. **(G)** Violin plots showing the expression of the top 50 DEGs (identified in the ‘single-PT’ setting) in the dominant pro-metastatic cells (Dominant) vs all the other cells (Other) in PT1 (upper panel) and PT5 divided in Group 1 and 2.

We then characterized upstream regulators of pro-metastatic genes, by enrichment analyses of promoter binding-motifs for known transcription factors (TFs), using the PASTAA tool [27]. Consistently with their functions, PT1 pro-metastatic genes were enriched for binding motifs of TFs implicated in the regulation of ECM remodeling/interaction (EGR1 and SP1 for ECM deposition; RUNX1 and MAZ for cell adhesion; MZF1 and FBI-1 for ECM degradation) and hypoxia response (SP1 and EGR1; Fig. 5E, upper panels). Likewise, PT5 pro-metastatic genes were enriched for binding motifs of TFs implicated in the regulation of dsRNA-IFN1 signaling (DEAF1 and IRF8) and UPR (HSF1 and PAX4; Fig. 5E, lower panels).

Together, these data indicate that both pro-metastatic clones overexpress genes involved in ECM interaction/remodeling, dsRNA-IFN1 signaling and stress response (hypoxia, unfolded protein, and antioxidant response). Notably, genes implicated in the same pathways are over-expressed in all other Group 2 clones as compared with Group 1 in most cases (Fig. 5C-D), suggesting that hyper-activation of these pathways is the distinguishing feature of pro-metastatic clones.

Finally, we investigated whether the identified pro-metastatic genes are homogeneously expressed across cells of the two pro-metastatic clones. For each clone, we assessed expression of the top 50 DEGs (‘*single-PT’* DE setting) at single-cell level. As shown in Fig. 5F, expression was highly heterogenous across pro-metastatic cells, with only a relatively-low cell fraction expressing >50% of DEGs above the corresponding PT mean (41 and 32% cells in PT1 and PT5 pro-metastatic clones, respectively), confirming that pro-metastatic clones retain high-degree of intra-clonal heterogeneity (see also Fig. S3B). Notably, pro-metastatic DEGs were also heterogeneously expressed by non-metastatic cells. Consistently with the enrichment of Group 2 cells in the two pro-metastatic clones, we observed a continuously decreasing expression-trend of pro-metastatic DEGs from pro-metastatic cells to Group 2 and Group 1 cells (Fig. 5G). Specifically, the fraction of cells expressing >50% DEGs was 21 and 1% in PT1 and PT5 non-metastatic Group 2 cells, and 18 and 4% in PT1 and PT5 non-metastatic Group 1 cells (Fig. 5F).

Thus, pro-metastatic clones are distinguished by the presence of a larger fraction of cells that simultaneously upregulate multiple genes involved in ECM interaction/remodeling, dsRNA-IFN1 response and stress response pathways, suggesting that concomitant hyper-activation of multiple pathways increases the likelihood of dissemination.

### Silencing of single pro-metastatic genes dramatically reduces metastatization *in vivo*

Finally, we investigated whether the pro-metastatic phenotype requires concomitant hyper-activation of multiple pathways, by silencing individual pro-metastatic genes *in vitro* and *in vivo*. We selected five targets (Fig. 6A): ANGPTL4, which is upregulated in both PT1 and PT5 pro-metastatic clones and involved in ECM degradation [28] and hypoxia response [29]; KCNQ1OT1, upregulated in the PT1 clone and involved in ECM degradation [30]; ITGB4, upregulated in the PT1 clone and involved in ECM adhesion [31]; LY6E and IFI6, upregulated in both pro-metastatic clones and involved in dsRNA-IFN1 signaling [32], [33]. Some of these genes have been previously implicated in the regulation of metastasis-associated phenotypes in other experimental contexts or cancer types, such as migration of BC cells (IFI6; [34], migration and liver colonization in melanoma (KCNQ1OT1; [35]), or metastatization in BC (ANGPLT4; [36]).

**Fig. 6.**
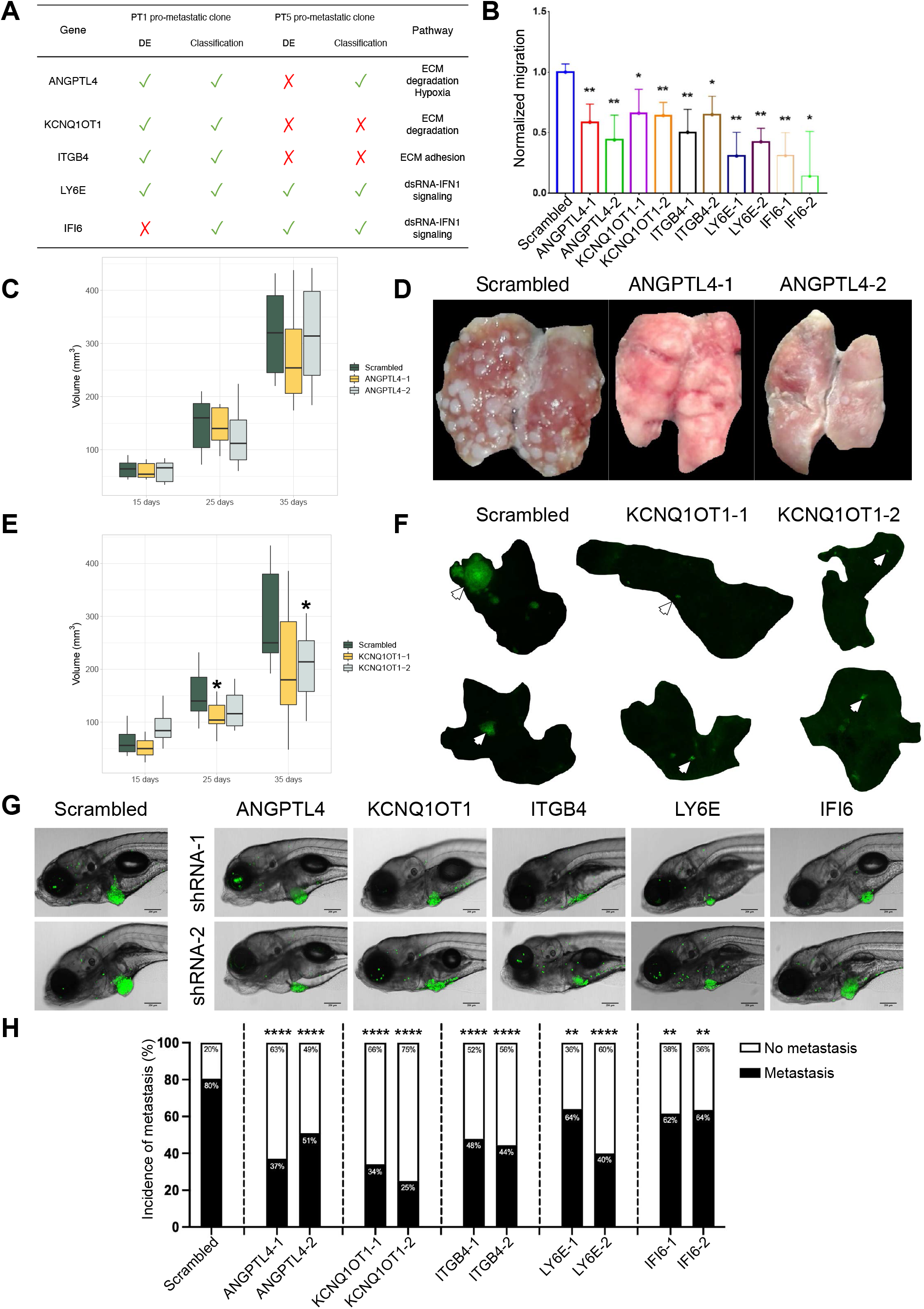
Silencing of single pro-metastatic genes dramatically reduces metastatization *in vivo*. **(A)** Schematic representation of the properties of the 5 pro-metastatic genes selected for *in vitro* and *in vivo* validation experiments. For each gene we report: the pro-metastatic clone where it was identified; the specific analysis (DE and/or classification) which allowed identification; and the functional pathway(s) in which is implicated. **(B)** Quantification of migration in the wound healing assay. Two images of the wound (at 0h and 8h) were used to compute migration as wound closure (see M&M). The wound closure in the shRNA populations of cells was normalized to the wound closure in the scrambled population (two-tailed T-test, * p < 0.05; ** p < 0.01; n ≥ 3). **(C)** Kinetic of PT growth *in vivo* with MDA-MB-231 infected with either the scrambled or the two ANGPTL4-shRNAs (Scrambled cohort n = 7; ANGPTL4_1 cohort n = 7; ANGPTL4_2 cohort n = 7). PT images were taken by IVIS-lumina *in vivo* system and PT volume was computed as (minimum diameter)^2^·(maximum diameter)/2. **(D)** Representative photos of lung metastases in mice carrying MDA-MB-231 cells infected with either the scrambled or the two ANGPTL4-shRNAs. **(F)** Kinetic of PT growth *in vivo* with MDA-MB-231 infected with either the scrambled or the two KCNQ1OT1-shRNAs (Scrambled cohort n = 7; KCNQ1OT1_1 cohort n = 7; KCNQ1OT1_2 cohort n = 7; two-tailed T-test, *p < 0.05). PT images were taken by IVIS-lumina *in vivo* system and PT volume was computed as (minimum diameter)^2^·(maximum diameter)/2. **(G)** Representative images of lung slices in mice carrying MDA-MB-231 cells infected with either the scrambled or the two KCNQ1OT1-shRNAs. MDA-MB-231 cells in the lung parenchyma can be distinguished because they carry the H2B-GFP fluorescent reporter. Metastatic nodules are indicated by arrows. **(H)** Representative images of zebrafish larvae apical portion (scale bar = 200 µm), showing the primary tumor mass at the site of PVS injection. MDA-MB-231 cells in the PVS can be distinguished because they carry the H2B-GFP fluorescent reporter. **(I)** Bar plots of metastasis incidence in zebrafish larvae computed 4 days after transplantation (scrambled cohort n = 434; ANGPTL4_1 cohort n = 59; ANGPTL4_2 cohort n = 48; KCNQ1OT1_1 cohort n = 47; KCNQ1OT1_2 cohort n = 27; ITGB4_1 cohort n = 46; ITGB4_2 cohort n = 37; KLY6E_1 cohort n = 50; LY6E_2 cohort n = 25; IFI6_1 cohort n = 66; IFI6_2 cohort n = 77; T-test for proportions, *p < 0.05; ** p < 0.01; *** p < 0.001; **** p < 0.0001). Metastasis incidence represents the number of fish displaying at least one metastasis divided by the total number of fish.

MDA-MB-231 cells were engineered to stably express the histone 2B-GFP (H2B-GFP) reporter and two independent short-hairpin RNAs (shRNAs) for each of the 5 genes. shRNAs-mediated silencing is shown in Fig. S6A. Silencing of ANGPTL4 and IFI6 did not affect *in vitro* growth of MDA-MB-231 cells, which was instead slightly, yet significantly, reduced by silencing of KCNQ1OT1, LY6E and ITGB4 (Fig. S6B). In no case we observed effects on cell-cycle distribution of MDA-MB-231 cells (Fig. S6C). For all targets, instead, silencing reduced significantly *in vitro* migration (from ∼40 to ∼80%), as determined by the wound-healing assay (Fig. 6B).

Effects on metastasis *in vivo* was assessed in mouse (ANGPTL4 and KCNQ1OT1) and zebrafish (all the 5 selected genes). *Per* each mouse, 200,000 MDA-MB-231 cells expressing scrambled, ANGPTL4- or KCNQ1OT1-shRNAs were injected orthotopically in the mammary gland. Silencing of ANGPLT4 exerted no significant effects on PT growth (Fig. 6C), while it dramatically reduced numbers of metastatic nodules in the lungs (Fig. 6D), as previously reported in MDA-MB-231 [36]. Silencing of KCNQ1OT1 slightly reduced PT growth (Fig. 6E) and decreased lung infiltration (Fig. 6F), with a ∼70% reduction of the metastatic burden (which quantifies the lung infiltration normalized on the primary mass; see Fig. S6D and Materials and Methods) with both shRNAs. ∼500 H2B-GFP-MDA-MB-231 cells expressing scrambled, ANGPTL4- or KCNQ1OT1-shRNAs were also microinjected in the Peri-Vitelline Space (PVS) of zebrafish *larvae* obtained at two days post-fertilization. *Larvae* were then incubated at 34°C and GFP-positive cancer-cells monitored by imaging of the transparent larval-body at 4 days post injection (dpi). Injected cells form a primary mass at the local site of injection, enter circulation, extravasate and seed at distant sites, where they eventually expand to metastatic nodules (> 5 cells). Silencing of ANGPTL4 and KCNQ1OT1 gave in zebrafish the same results observed in mice: no (ANGPTL4-shRNAs) or significant (KCNQ1OT1-shRNAs) effects on PT growth (Fig.6G for representative images and S6F for quantitative analyses) and dramatic reduction of metastasis incidence for both (ANGPTL4- and KCNQ1OT1-shRNAs; Fig. 6H). The same experiment was performed for the other three pro-metastatic genes (IFI6, ITGB4 and LY6E). IFI6-shRNAs did not affect PT growth, which was instead significantly reduced by ITGB4- or LY6E-shRNAs (Fig. 6G and S6F). Notably, silencing of these three targets significantly reduced metastasis incidence (Fig. 6H).

Together, these data demonstrate that acquisition of the pro-metastatic phenotype requires concomitant expression of individual genes involved ECM remodeling/interaction, dsRNA-IFN1 signaling and hypoxia response. Our data also revealed a previously undescribed role in the metastatic progression of BC of new targets, such as KCNQ1OT1, ITGB4, LY6E and IFI6. Notably, these pro-metastatic genes are not necessarily critical for tumor growth, while they are indispensable for the ability of BC cells to migrate *in vitro*.

## Discussion

We report here the first characterization of the transcriptional determinants of the pro-metastatic phenotype in BC at the single-cell level, and showed that they involve concomitant hyper-activation of ECM remodeling/interaction, dsRNA-IFN1 signaling and different branches of the integrated stress response (i.e., hypoxia, UPR and antioxidant response). Notably, the identified pro-metastatic transcriptional determinants contributed to the genesis of gene-expression signatures that function as strong predictors of BC patient metastatic relapse.

Each of these pathways has previously been implicated in metastasis formation in several tumor-types, including breast cancer [4] ECM remodeling/interaction activates migration and local invasion, which are obligatory early-steps of the metastatic cascade [37]. Though UPR, hypoxia and IFN1 signaling are potent inducers of apoptosis [38], [39], [40], their persistent activation promotes cancer cell survival, local invasion and metastasis formation [18], [41], [42]. Coherently, in our data, we did not observe up-regulation of known mediators of apoptosis induced by UPR (e.g., NOXA, BIM, PUMA or DR5) [43], hypoxia (e.g., HIF-1α or p53) or dsRNA-IFN1 signaling (e.g., the 2-5A-dependent ribonuclease RNASEL, components of the death-receptor signaling pathways or of the caspase cascade [44]. We found, instead, moderate activation of several pro-survival transcriptional targets of both pathways, including chaperones and antioxidant response genes. Thus, our data strongly indicate that pro-metastatic BC cells mount pro-survival adaptive stress response pathways to cope with stress signals from the microenvironment, such as hypoxia, or from inside cancer cells, such as dsRNA from active endogenous retroelements [45] or hyper-proliferation-induced endoplasmic reticulum stress [46].

Activation of these pathways occurs, however, in most other PT cellular-clones, suggesting that it is not sufficient *per se* to initiate the metastatic process. Indeed, our analyses revealed that the same pathways are heterogeneously activated across all PT clones along a gradient that inversely correlates with clonal proliferative potential. Widespread intra-clonal transcriptional heterogeneity has already been described in a murine BC model analyzed by multicolor lineage tracing, where most of the identified clones contained hyper-proliferative cells or cells that had either undergone EMT [47].

Rather, the distinctive traits of pro-metastatic clones are the concomitant activation of multiple pathways, and, for each pathway, the overexpression of specific genes in a higher fraction of cells (pathways hyper-activation). In particular, pro-metastatic clones overexpress genes involved in ECM remodeling/interaction, dsRNA-IFN1 signaling, hypoxia, and UPR genes involved in the regulation of protein homeostasis (HSPA1A (Hsp70.1), HSP90AA1, DNAJB1 (Hsp40), and HSPD1 (Hsp60)). Notably, we also observed up-regulation of several antioxidant genes (TXNRD1, MGST1, PTGES (MGST1-L1)) known to be critical for the survival of cancer cells at multiple stages of the metastatic process [9].

Strikingly, we showed that the inactivation of individual genes involved in either ECM remodeling/interaction or dsRNA-IFN1 signaling is sufficient to hinder metastasis spreading, demonstrating that simultaneous activation of multiple pathways is critical to initiate the metastatic cascade. Consistently, we observed higher fraction of cells in the pro-metastatic clones that simultaneously express pro-metastatic genes of different signaling pathways, as compared to the non-metastatic clones. Together, these observations suggest that crosstalk among different pathways plays a major role in the initiation of the metastatic cascade. Notably, dsRNA-IFN1 was recently implicated in the activation of invasive phenotypes through genes that are upregulated in our pro-metastatic clones (i.e., IFI6 [34], IFI27 [48], IFITM1 [49], or ISG15 [50]. Similarly, UPR activates migration/invasion through the release of the CXCL2, 3, 8 proinflammatory chemokines (which induce lamellipodia and filopodia formation [51], [52], uPAR (the receptor of urokinase-type plasminogen activator, which mediates ECM degradation [52], and UPR-induced antioxidant response genes [53], [54]. In summary, concomitant hyper-activation of pro-survival UPR signaling might support protein homeostasis leading to persistent antioxidant response, which synergizes with sustained hypoxia response and dsRNA-IFN1 signaling to activate ECM remodeling/interaction and, ultimately, the invasive phenotype.

Similar to previous results with genetic barcoding of MDA-MB-231 cells [20] and TNBC PDXs [21], [22], we showed that clonal complexity in metastases is significantly lower than in PTs. Our data suggest that only a few clones in the PT acquire metastatic potential, despite most clones possess the potential to generate the same adaptive response phenotypes. In this respect, we showed that the transcriptional profile is significantly lineage-dependent. However, the metastatic potential of clones is largely independent of the lineage, therefore indicating a major role for micro-environmental or inside-cell signals in promoting metastatization. Importantly, this scenario contrasts with previous findings in a lung cancer model, where the dissemination potential was found to be ingrained in the clonal identity [55]. Alternatively, both lineage and environmental context can influence the expression of the pro-metastatic phenotype, with the latter determining the metastatic phenotype only within a subset of lineage-restricted cells. The limited number of pro-metastatic clones analyzed so far, however, represents a major limit of these analyses.

We found that metastases are almost monoclonal, with a single clone accounting for >90% cells. Our finding significantly differs from what reported in MDA-MB-231 cells using either genetic [20] or multicolor barcoding [56]. However, primary breast tumors were obtained in the former study by subcutaneous heterotopic transplantation, while in the latter the PT size at resection was much smaller (100 mm^3^ vs 500-1000 mm^3^ in our work) and polyclonal metastasis coexisted with monoclonal metastasis at other anatomical sites. These last results suggest that specific environmental contexts at the PT or distant sites may influence the capacity of single disseminated clones to expand upon seeding. Identification of the environmental factors in the primary tumor or at distant sites that influence the growth potential of disseminated clones is of obvious clinical relevance.

Finally, consistently with previous reports [21], [22], [57], we showed that pro-metastatic clones are highly under-represented in the primary tumor. Analyses of inter-clonal transcriptional heterogeneity showed an inverse correlation between numbers of proliferating cells and activation of pro-metastatic stress-response pathways (i.e., hypoxia, dsRNA accumulation, and UPR), which are known to induce transient cell-cycle arrest [57], [58], [59], [60]. Single clones which experience this conditioning thus remain under-represented in the PT and hyper-activate specific response pathways (i.e., dsRNA-IFN1 signaling and ECM remodeling/interaction) that are permissive for metastatization. Notably, the low fitness of pro-metastatic clones is reversible upon seeding at distant sites.

In conclusion, our work revealed that the pro-metastatic phenotype in BC mainly arises in rare PT clones in a lineage-independent manner and is characterized by the concomitant hyper-activation of genes involved in ECM remodeling/interaction, dsRNA-IFN1 signaling and stress responses. Pathway hyperactivation correlates with a temporary growth arrest of the pro-metastatic clones in the PT, which is rapidly reversed upon distant organ seeding.

## Authors contributions

Conceptualization, N.R., A.C., and P.G.P.; Software, A.C., R.H., A.T., S.C., and E.G.; Formal analysis, N.R., A.C., R.H., A.T., and S.C.; Investigation N.R., A.C., F.R., C.P., A.D., V.G., G.B., A.P., and P.F.; Writing - Original Draft, N.R., A.C., and P.G.P.; Visualization, N.R., A.C., and P.G.P.; Supervision, E.G., L.M., L.L., E.M., and P.G.P.; Project administration, P.G.P.; Funding acquisition, P.G.P.

## Acknowledgements

We thank Luisa Lanfrancone, Gioacchino Natoli, and Ruggero De Maria for many critical discussions; the members of the IEO Genomics, FACS and sorting, Tissue Culture, and Imaging facilities for technical support. We thank the members of the Cogentech animal facility for the mouse and zebrafish care. We are grateful to Martin Schaefer, Tommaso Leonardi, Stefania Averaimo, Camilla Cerutti, Fabio Alfieri, Danilo Cagnina, Camilla Ugolini, and Lorenzo Ciacci for fruitful discussion and for their valuable technical support. This work was supported by PRIN (Ministry of University and Research) n. 2017L8FWY8. N.R. was supported by is supported by a fellowship from Fondazione IEO-CCM.

## Materials & Methods

## CONTACT FOR REAGENT AND RESOURCE SHARING

Further information and requests for resources and reagents should be directed to and will be fulfilled by the Lead Contact, Pier Giuseppe Pelicci

### Materials availability

This study did not generate new unique reagents.

### Data and code availability

Single-cell RNA-seq data will be deposited at European Nucleotide Archive (ENA) and publicly available as of the date of publication. Microscopy data reported in this paper will be shared by the lead contact upon request. All original code will be deposited and publicly available as of the date of publication. Any additional information required to reanalyze the data reported in this paper is available from the lead contact upon request.

## EXPERIMENTAL MODEL AND SUBJECT DETAILS

### Cell lines

All cells were cultured in adhesion in a 20% O2, 5% CO2 incubator at 37°C. HEK293T and the metastatic human TNBC cell line MDA-MB-231 were purchased from the American Type culture collection (ATCC) and cultured in DMEM (EuroClone), supplemented with 10% South American fetal bovine serum (FBS; Microtech), 2mM L-glutamine (EuroClone), and 100U/ml penicillin-streptomycin (Thermo Fisher Scientific). All the cell lines were tested for mycoplasma contamination routinely. All the cell lines were split once they reached ∼80% confluence and cultured in vitro for no more than 10 passages after thawing.

### Mice

Female NOD/SCID Il2-Rγ null (NSG) mice were purchased from Charles River Laboratory and housed under specific pathogen-free conditions at 22 ± 2°C with 55 ± 10% relative humidity and with 12 h day/light cycles in mouse facilities at the European Institute of Oncology–Italian Foundation for Cancer Research (FIRC) Institute of Molecular Oncology (IEO–IFOM, Milan, Italy) campus. *In vivo* studies were performed after approval from our fully authorized animal facility and our institutional welfare committee (OPBA) and notification of the experiments to the Ministry of Health (as required by the Italian Law (D.L.vo 26/14 and following amendments); IACUCs Numbers: 833/2018, 679/2020), in accordance with EU directive 2010/63.

### Zebrafish

Zebrafish (Danio rerio) from *Tg(kdrl:DsRed)* strain were housed in compliance with national guidelines regarding animal welfare (as required by the Italian Law (D.L.vo 26/14 and following amendments)). Fish were housed in a “Aquatic Habitats” husbandry system, in 3l water tanks with a density of 0.2l per fish at 28.5± 2°C, water pH=7 and conductivity=500 µS. The circadian rhythm imposed was 14 hours of light and 10 hours of darkness. All the xenotransplantation and imaging experiments presented in this study were performed on animals at larval stage. Zebrafish larvae (from 2 dpf to 6 dpf) were raised in Petri dishes filled with E3 water medium (5 mM NaCl, 0.17 mM KCl, 0.33 mM CaCl, 0.33 mM MgSO4 and 0.05% methylene blue in ddH2O), at a density of ∼60 fish per plate. 18-22 hours post fertilization larvae were transferred to E3 water supplemented with 0.003% N-phenylthiourea (PTU, Sigma-Aldrich) to inhibit melanin formation and conserve optical transparency.

## METHODS

### Plasmid production

The Perturb-seq GBC library (pBA571) [23], [24] was purchased from Addgene and used for lineage tracing experiments. This vector contains a random 18-nt guide barcode (GBC) between the blue fluorescent protein (TagBFP) and polyadenylation signal sequences. The GBC was inserted by Gibson assembly using subsequently disrupted EcoRI and AvrII sites. The vector contains puromycin and ampicillin resistance genes and the reporter gene TagBFP constitutively expressed under the control of EF1α promoter. We estimated the complexity of this library by two subsequent rounds of sequencing (30 million reads per sequencing run). In the first round, we identified 3,152,856 unique GBCs (median reads per unique GBC = 1), while in the second round we identified 2,386,607 unique GBCs (median reads per unique GBC = 1). The overlap between the two rounds was of 797,353 GBCs, which represented the 33.41% of the unique GBCs in the first round. Therefore, we reasoned that the initially sequenced 3,152,856 unique GBCs represented the 33.41% of the whole library, whose complexity was estimated to range between 5 and 10 million unique GBCs. To amplify the Perturb-Seq GBC library, we transformed *E. coli* ElectroMAX™ DH5α-E™ electro-competent cells (Thermo Fisher Scientific), as indicated by the authors of the library [23], [24]. Briefly, we added the DNA to microcentrifuge tubes, thawed cells on ice, and distributed 20µl of bacteria in the chilled microcentrifuge tube. We pipetted the cell/DNA mixture into a chilled 0.1 cm cuvette and electroporated with BioRad GenePulser® II electroporator (2.0 kV, 200 Ω, 25 μF). Bacteria were let to recover in 1 ml of S.O.C. medium (Thermo Fisher Scientific) for 1 hour at 37°C (600rpm). Then, 5μl of recovered cell suspension were plated on selection plates (Agar + 50 mg/L ampicillin (Sigma-Aldrich)) to assess transformation efficiency. The rest of recovery was grown overnight in 500 mL of Luria Broth + ampicillin while shaking at 37°C. The day after, we performed DNA extraction with NucleoBond Xtra Maxi (Macherey-Nagel) kit according to the volume of growth and the manufacturer’s instructions.

The pRSI17-U6-sh-UbiC-TagGFP-2A-Puro Plasmid was purchased from Cellecta and used in target validation experiments. This vector allows for the constitutive expression of a short hairpin RNA (shRNA) under the U6 promoter. The vector contains puromycin and ampicillin resistance genes and the reporter gene green fluorescent protein (TagGFP) constitutively expressed under the UbiC promoter. shRNA sequences are reported in the **Suppl. Table 9**. The shRNA cloning was performed according to manufacturer instructions. Briefly, sense and antisense oligos were diluted to a final concentration of 20μM and mixed in a 1:1 ratio. Phosphorylation and annealing reactions were performed in 20μl of 1X T4 Polynucleotide Kinase Reaction Buffer (New England Biolabs). 10 units of T4 Polynucleotide Kinase (New England Biolabs) were added to the mix and the solution was incubated for 30 minutes at 37°C to allow phosphorylation. Then, the reaction was heated to 95°C for 2 minutes to disrupt secondary structures within oligonucleotides. Ultimately, the mix was cooled down gradually to room temperature to allow for sense and antisense strands to anneal. Annealed oligonucleotides were diluted 1:10 in 1X T4 Polynucleotide Kinase Reaction Buffer and ligation was performed in 10μl of 1X T4 DNA Ligase Buffer (New England Biolabs). 1μl of annealed and diluted oligonucleotides were mixed with 40 units of T4 DNA Ligase (New England Biolabs) and 1μl of pRSI17-U6-sh-UbiC-TagGFP-2A-Puro Plasmid. The reaction was stored at room temperature for 4 hours protected from light. After ligation, the reaction was used to transform Stbl3 bacteria by heat shock. Ultimately, 100µl of cell suspension were plated on selection plate (Agar + 50 mg/L ampicillin (Sigma-Aldrich)) and incubated overnight at 37°C. The day after, 3 colonies were picked and grown overnight, to perform DNA extraction. For DNA extraction, we used NucleoBond Xtra Maxi (Macherey-Nagel) kit according to the volume of growth and the manufacturer’s instructions. Plasmids were Sanger-sequenced to check the quality of both ligation and sequence.

The pLenti CMV Puro LUC (w168-1) was purchased from Addgene and used in in vivo experiments with NSG mice. This 3rd generation lentiviral vector allows the constitutive expression of firefly luciferase under the CMV promoter. The vector contains genes that encode for puromycin and ampicillin resistance.

The Tet-Off-H2B-GFP lentiviral vector was kindly provided by F. Ruscitto (Department of Experimental Oncology, European Institute of Oncology, Milan) [61] and used in target validation experiments. This 3rd generation lentiviral vector allows the constitutive expression of the fusion protein histone H2B-GFP under the control of a tetracycline-responsive promoter element (TRE), which is in turn regulated by a tetracycline-controlled transactivator protein (tTA).

### Viral production

Lentiviruses were packaged in HEK293T cells using pMDLg/RRE, pRSV-REV, and pMD2.G third generation packaging and envelope plasmids. pRSI17-U6-sh-UbiC-TagGFP-2A-Puro, pLenti CMV Puro LUC (w168-1), and Tet-Off-H2B-GFP lentiviral vectors were transfected by calcium phosphate [62]. Briefly, 10μg of transfer plasmids were mixed with 4μg of pMD2.G, 3μg of pRSV-REV, and 4μg of pMDLg/pRRE packaging vectors per each 10cm plate of HEK293T. CaCl2 (Sigma-Aldrich) is added to the vectors mix to a final concentration of 0.5 M. The mix was added drop-wise to 500 μl of 2X HBS (HEPES buffered saline (Thermo Fisher Scientific): 250 mM HEPES pH 7.0, 250 mM NaCl and 150 mM Na2HPO4), constantly bubbling. Chloroquine (Sigma-Aldrich) was added to HEK293T medium to the final concentration of 100μM to increase transfection efficiency. After 15 minutes of incubation at room temperature, the precipitate was added drop-wise to the plates and immediately mixed with the culture medium by gentle swirling.

To transfect the Perturb-seq GBC library we performed Lipofectamine™ (Thermo Fisher Scientific) transfection as indicated by the authors [23], [24]. Briefly, per each 15cm plate of HEK293T, solution A (containing 25μg of Perturb-seq GBC library, 10μg of pMD2.G, 7.5 μg of pRSV-REV, 10μg of pMDLg/pRRE, 60μl of Plus™ Reagent in 1.2 ml of DMEM without FBS and antibiotics) and solution B (containing 90μl of Lipofectamine™ in 1.2 ml of DMEM without FBS and antibiotics) were incubated at RT for 15 minutes and then mixed together. After a further 15-minute incubation, the DNA-lipid complexes were slowly added to the HEK293T cell monolayer.

Viral supernatants were collected 2-4 d post-transfection and filtered through 0.45 mm filters. Filtered supernatants were concentrated by ultracentrifugation at 22,000 rpm for 2 hours at 4°C with OptimaTM L-90K Ultracentrifuge (Beckman Coulter) and stored at -80°C (never refrozen).

### *In vivo* lineage tracing experiments

To perform *in vivo* lineage tracing experiments, MDA-MB-231 cells were transduced with the Perturb-seq library by a single round of infection on adherent cells. Cells were plated in 10cm plates at a density of 2·10^6^ cells/plate and infected with multiplicity of infection (MOI) = 0.5 in the presence of 8μg/ml of Polybrene (Sigma-Aldrich). Transduction efficiency was measured by FACS analysis, which revealed that 30% of cells were successfully infected: assuming a Poisson distribution for the number of viral integrations per cell, this percentage of infected cells guarantees that ∼90% of cells are infected by one single viral particle.

Infected cells were selected by puromycin (2.5μg/ml for 72h; Vinci-Biochem) and allowed to expand for additional 72h at the end of the selection. Ultimately, cells were transduced with pLenti CMV Puro LUC (w168-1) reporter vector at MOI = 5 to ensure that most cells expressed luciferase. Prior to *in vivo* transplantation, ∼10^6^ cells were set apart to assess the distribution of clones in the *in vitro* reference population (as discussed in detail below). Cells from two independent *in vitro* reference populations were transplanted in 6 NSG mice (3 mice per each reference). Per each mouse, 200,000 MDA-MB-231 cells were resuspended in 30μl phosphate buffered saline (PBS (Thermo Fisher Scientific)) and growth-factor-reduced Matrigel (Corning) in a ratio of 1:1 and then injected in the ninth mammary gland by intra-ductal transplantation without prior clearing of the fat pad. Tumor growth was weekly monitored by IVIS-Lumina *in vivo* imaging system (Perkin Elmer) and tumor volume estimated by the formula (minimum diameter)^2^·(maximum diameter)/2 [63].

Tumors were resected ∼40 days post injection (once they reached a volume of 0.5-1 cm^3^). Immediately after resection, MDA-MB-231 PTs were minced in small pieces and single-cell suspensions prepared by digestion for 1h at 37°C with 200 U/mL collagenase (Sigma-Aldrich), 100 µg/mL hyaluronidase (Sigma-Aldrich), and 100U/ml pen-strep (Thermo Fisher Scientific) in DMEM (Euroclone). After digestion, ∼10^6^ cells were set apart to assess the distribution of clones in the PTs (as discussed in detail below). The remaining PT cells were frozen in 90% South American FBS (Microtech) + 10% Dimethyl Sulfoxide (Sigma-Aldrich) and stored at - 80°C.

Upon resection, lung infiltration was weekly monitored by IVIS-Lumina *in vivo* imaging system and mice sacrificed 21 days after PT resection. Lungs and liver were collected, and single-cell suspensions prepared. To isolate metastatic cells from liver and lungs, the organs were minced in small pieces and single-cell suspensions prepared following the Miltenyi Tumor dissociation kit (Miltenyi, https://www.miltenyibiotec.com/upload/assets/IM0002061.PDF). After digestion, suspensions were frozen in 90% South American FBS (Microtech) + 10% Dimethyl Sulfoxide (Sigma-Aldrich) and stored at -80°C. The percentage of alive and GBC-barcoded cells in both PTs and metastases was assessed by flow cytometry on freshly thawed samples.

### Immunohistochemistry

One MDA-MB-231 PT was cut into pieces upon resection to perform immunohistochemistry (IHC). The sample was fixed for 24h in 4% formaldehyde (Sigma-Aldrich) and subsequently moved to 70% ethanol (Panreac Applichem) prior to paraffin inclusion. Proliferating cells were identified by staining with the rabbit monoclonal anti-human Ki67 (SP6 ab16667, Abcam) antibody (5μg/ml) and revealed by the immunoreaction product brown diaminobenzidine (DAB). Ki67+ cells were quantified using the ImageJ plugin ImmunoRatio [64], through calculation of DAB-stained area/total nuclear area.

### DNA extraction for bulk targeted sequencing

To measure *in vitro* and *in vivo* clonal distributions, we extracted DNA from MDA-MB-231 cells. For each sample, ∼10^6^ cells were combined with a mixed population of spike-in controls. Cells were lysed in TrisHCl pH 8.0 (VWR) 10mM, EDTA (VWR) 0.5mM, SDS (Sigma-Aldrich) 0.2%, NaCl (Sigma-Aldrich) 100mM and proteinase K (Sigma-Aldrich) 0.1mg/ml for 4h at 56°C. Proteinase K was then inactivated by incubating the suspension at 80°C for 10 minutes. Upon lysis, DNA was precipitated by the addition of 1 volume of cold isopropanol (Panreac Applichem) and subsequently washed twice in 70% ethanol (Panreac Applichem) prior to the resuspension in water.

### Targeted sequencing library preparation

PCR of DNA-integrated barcodes was performed on 200ng of genomic DNA using the following PCR setting (primers are listed in **Suppl. Table 9**):

| Process | Temperature (°C) | Duration (s) | Cycles |
| --- | --- | --- | --- |
| Initial denaturation | 98 | 30 | 1 |
| Denaturation | 98 | 7 | 25 |
| Annealing | 62 | 25 |  |
| Extension | 72 | 20 |  |
| Final extension | 72 | 60 | 1 |

The presence of the diagnostic 240bp PCR-product was checked for every sample using 2% agarose gel electrophoresis. PCR products were purified by QIAquick PCR purification kit (Qiagen) and 2ng per sample sequenced on NovaSeq 6000 Sequencing System (Illumina), with a depth of sequencing equal to 30 million reads/sample.

### Spike-in controls preparation and sequencing

To estimate the size of any MDA-MB-231 clone (i.e., its absolute cell number) from the number of reads containing its GBC, we generated a set of five spike-in control populations (i.e., cellular populations of known size, each marked with a different GBC; list of spike-in controls in **Suppl. Table 9**) as in [20] Since we did not need a high coverage of GBC library complexity to generated spike-in controls, Stbl3 bacteria were heat-shock-transformed, and 5 independent colonies were picked and grown overnight. Plasmid DNA was extracted with NucleoBond Xtra Maxi (Macherey-Nagel) kit and Sanger-sequenced.

The Spike-in control plasmids were used to produce lentivirus as previously described. Then, we infected 5 independent MDA-MB-231 populations and selected cells by puromycin administration for 72 hours. By serial dilution, we prepared 5 spike-in control populations with increasing size (100 - 1,000 - 10,000 - 100,000 - 1,000,000 cells respectively). Spike-in controls were mixed together and stored at -20°C. Spike-in controls were combined with our samples of interest (*in vitro* references and PTs) prior to DNA extraction for sequencing.

### Targeted sequencing reads pre-processing

.*fastq* files from targeted sequencing and single-cell sequencing (see below) were pre-processed with a custom nextflow pipeline similar to [24] Each sample files were pre-processed separately. Pre-processing of single-cell sequencing reads from a sample (see below) was performed upon pre-processing of targeted sequencing reads from its matched reference sample. Considering targeted sequencing reads, GBC-containing sequences were extracted searching for their characteristic upstream ‘anchor’ sequence (i.e., GGGTTTAAACGGGCCCTCTAG) with 1 mismatch tolerance. Then, sequences with length <18 bp were filtered out, and whitelisted with umi-tools [65] to create a whitelist of putatively correct GBC sequences. This whitelist was used to correct all GBC sequences in targeted sequencing reads, whose counts were then used as input for clonal distribution analysis.

### Clonal distribution analysis

We first estimated the linear relationship between GBC read counts and populations sizes by simple linear regression, taking advantage of spike-ins data. This model was then used as a titration curve to estimate the (unknown) sizes of other putative clones from their GBC read counts. We excluded GBCs whose estimated clone size felt outside the titration curve range [100, 10^6^ cells]. This step excluded ∼3-9% of the total GBC-containing reads per sample, which is in line with previous studies [21], [22]. Retained GBCs and related cellular populations were considered as ‘clones’. For each sample, clones’ frequencies were computed as clones’ sizes over the total estimated number of sample cells. To quantify the clonal complexity of a sample, we calculated the Shannon Entropy (SE) of its clonal distribution:

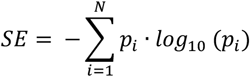

where pi is the clonal frequency of clone *i*, with *i* = 1, …, *N* and *N* is the sample clone number.

### Single-cell library preparation and sequencing

Cancer cells were isolated from PTs and metastases via fluorescence activated cell sorter (FACS) using a BD FACSAria II. After gating for singlets and live cells (alive cells were identified by propidium iodide (250µg/ml; Sigma-Aldrich) negativity), TagBFP-positive (GBC-barcoded) cells were sorted. For samples with a high yield of cells (PTs) 100,000 cells were sorted on the purity setting. For each of the lung and liver samples, the entire sample was sorted on the purity setting to recover as many TagBFP-positive cells as possible. After sorting, samples were passed through a 30 mm filter and then centrifuged at 1300rpm for 5 min. Supernatant was removed and cells resuspended to a final concentration of 500-1,000 cells/µl in PBS (Thermo Fisher Scientific). The single-cell suspension was mixed with reverse transcription mix using the Chromium Single-cell 3’ reagent kit protocol V3.1 (10x Genomics) and loaded onto 10x Genomics Single Cell 3’Chips to generate single-cell beads in emulsion. The gel beads are coated with unique primers bearing 10× cell barcodes, unique molecular identifiers (UMI) and poly(dT) sequences. The chip was then loaded onto a Chromium instrument where RNA transcripts from single cells are reverse-transcribed. cDNA molecules from one sample were pooled, amplified, and the amplified cDNAs were fragmented. Final libraries were generated by incorporating adapters and sample indices compatible with Illumina sequencing and quantified Qubit (ThermoFisher Scientific). The size profiles of the sequencing libraries were examined by Agilent Bioanalyzer 2100 using a High Sensitivity DNA chip (Agilent). Two indexed libraries were equimolarly pooled and sequenced on Illumina NOVAseq 6000 Sequencing System using the v1.5 Kit (Illumina) with a customized paired end, dual indexing (26/8/0/91-bp) format according to the manufacturer’s protocol. A coverage of ∼50.000 reads per cell was adopted for each sequencing run.

In addition, upon cDNA preparation, as the number of GBC-supporting reads per cell was low [23], [24], we performed a semi-nested enrichment PCR to increase the number of GBC-supporting reads. In the 1st reaction, we amplified 5ng of starting cDNA material and used the following setting (primer sequences are reported in **Suppl. Table 9**):

| Process | Temperature (°C) | Duration (s) | Cycles |
| --- | --- | --- | --- |
| Initial denaturation | 98 | 10 | 1 |
| Denaturation | 98 | 5 | 22 |
| Annealing | 72 | 5 |  |
| Extension | 72 | 10 |  |
| Final extension | 72 | 60 | 1 |

The presence of the diagnostic ∼400bp product was checked for each sample using 2% agarose (Canvax Biotech) gel electrophoresis. The PCR product was diluted 1:1000 to reduce the amount of aspecific products.

Then, we performed a 2nd reaction with the following primers and setting (primer sequences are reported in **Suppl. Table 9**):

| Process | Temperature (°C) | Duration (s) | Cycles |
| --- | --- | --- | --- |
| Initial denaturation | 98 | 10 | 1 |
| Denaturation | 98 | 5 | 20 |
| Annealing | 62 | 5 |  |
| Extension | 72 | 10 |  |
| Final extension | 72 | 60 | 1 |

The presence of the diagnostic ∼400bp product was checked for every sample using 2% agarose (Canvax Biotech) gel electrophoresis. PCR products were purified with QIAquick PCR purification kit (Qiagen) and 2ng per sample sequenced on NovaSeq 6000 Sequencing System (Illumina), with a depth of sequencing equal to 30 million reads/sample.

### Single-cell sequencing reads pre-processing

Single-cell paired-end reads were pre-processed as follows (R1 and R2 refers to read1 and read2 sequence records, respectively). First, trimmed (first 33 nt) R2 sequences were aligned with bowtie2 [66] (1 mismatch tolerance) to an index built from the TAGCAAACTGGGGCACAAGCTTAATTAAGAATT sequence (the R2 GBC identifier). Then, aligned read names were used to split R1 and R2 .fastq files into ‘GBC-containing’ and ‘transcriptional’ files with seqtk [67] subseq. 10x Cellular barcode (CBC) (R1), UMI (R1) and GBC sequences (R2) were subsequently extracted from each GBC-containing read pair. In parallel, CBC sequences (first 16 bp of each R1 sequence) were extracted from transcriptional R1 sequences, whitelisted with umi-tools whitelist method, and corrected as previously described (see above). At this point, bowtie2 was used to align GBC sequences extracted from single-cell GBC-containing reads to an index built from the GBC sequences whitelisted in the pre-processing step of the matched *in-vitro* targeted sequencing reference sample. Only aligned GBC sequences were subsequently matched to their previously extracted CBC and UMI, and used to retrieve and count unique CBC-GBC combinations. In parallel, transcriptional reads and whitelisted CBCs were used to build a filtered sample gene expression (GE) matrix with STAR-solo [68] Each sample GE matrix and matched CBC-GBC table (storing unique CBC-GBC combinations counts) was used as an input for Cell and Gene Quality control step.

### Cell and gene quality control

Cell and gene quality control (QC) were performed for each sample separately.

First, putative cells (CBCs) assigned to multiple GBC or weakly assigned to a single GBC (CBC-GBC combination supported by <= 5 UMIs or <= 30 reads) were filtered out as in [24] Then, genes expressed by less the 5 cells and cells with QC metrics (i.e., number of UMIs and number of genes detected per cell) falling outside of the [median – 5*M.A.D., median + 5*M.A.D.] range were discarded. Additionally, cells with cell complexity > 0.8 and mitochondrial genes percentage > 50% (as adopted by [69]. for this cell line) was filtered out. Finally, the scDblFinder [70] method was run to filter out other putative cell doublets.

### Gene expression matrices processing, clustering and visualization

Each sample cleaned GE matrix was log-normalized with the scran method [71], [72] After normalization, GE matrices were subsetted for common genes and concatenated together.

To obtain a robust set of hypervariable genes (HVGs), first the gene-wise the mean-variance trend was modelled for each sample using the modelGeneVar() function from scran. Second, the top 5000 genes ranked for their biological variation component was selected for each sample. Then, from the union of each of these sets, the top 2000 occuring genes were selected (breaking ties using their ranking according to their biological variation component). These HVGs were used to subset the concatenated log-normalized matrix. Unwanted technical (number of UMIs and mitochondrial percentage) and biological (only in the alternate ‘no cc’ workflow) sources of variation were regressed out from log-normalized expression values with the sc.tl.regressOut() function from scanpy [72]. The dimensionality of this corrected representation of gene expression data was then reduced with Principal Component Analysis (PCA). Batch (i.e., sequencing run) correction was performed on these data projections with the Harmony algorithm [73]. The resulting cell embeddings (first 30 components, henceforth referred to as ‘PCA space’) were finally used to construct the Shared Nearest-Neighbors (SNN) graph (constructed from an initial k-NN search with k=30, as implemented in the buildSNNGraph() function from scran) that fed Louvain clustering (implemented in the cluster_louvain() function from igraph [74]. We chose the clustering solution upon inspection of other possible clustering solutions, each obtained from the Louvain partitioning of different SNN graphs (each constructed from different initial k-NN searches, with k in [5,10,15,20,30,50]. For each clustering solution, we evaluated: i) clusters compactness and separation, with the Davies-Boudain (DB) index [75]. The DB index is calculated as the average similarity between each cluster *C*_*i*_ and its most similar one *C*_*j*_ for *i* = 1, …, *n* clusters. Specifically, similarity among two clusters *ij, R*_*ij*_ is defined as:

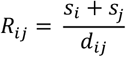

with *s*_*i*_ (*s*_*j*_) defined as the average distance between each point of cluster *i*, (*j*) and the centroid of that cluster, *C*_*i*_ (*C*_*j*_). The DB index is then defined as:

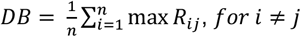

and ii) median kNN purity calculated as the median (over all cells in the dataset) of the quantity:

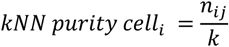

where the kNN purity of *cell*_*i*_ with membership in cluster *j* expresses the fraction of *cell*_*i*_ neighbors with membership in cluster *j, n*_*i,j*_, for a given neighborhood size, k; and iii) clusters biological annotation (derived from clusters markers and related over-represented pathways, see below). We chose the solution which best satisfied these notions of computational and biological separation. The PCA space were further embedded in 2 dimensions with Uniform Manifold Approximation and Projection (UMAP) [76] following scanpy workflow (i.e., sc.tl.neighbors() (k=30) and sc.tl.umap()(a=xy, b=xy)).

### Marker genes and cell cycle scoring

For each cluster, two markers list have been computed with the findMarkers() function from scran, using Wilcoxon test as test type. Only up-regulated genes (log2FC>0) with FDR>0.1 have been considered for subsequent analysis. Specifically, *‘all’* markers (i.e., genes found differentially expressed between a cluster and all the other clusters) have been used individually for cluster annotation, while *‘some’* markers (i.e., genes found differentially expressed between a cluster and at least 50% of all the others) have been used for Over Representation Analysis (ORA).

Assignment of each cell to a cell cycle phase was performed using the cellCycleScoring() function in Seurat [77].

### Differential Expression

All Differential Expression (DE) analyses were performed with the edgeR method [78], considering sequencing run (if necessary, considering the particular contrast), and gene detection probability as additional model covariates for each pairwise contrast, as suggested in [79]. Only up-regulated genes (log2FC>0) with FDR>0.1 have been considered for ORA. For the *‘Single-NN’* setting, DE was performed between either PT1 or PT5 pro-metastatic cells and their ‘neighboring’ non-metastatic cells, within the same PT. For each of these two contrasts, neighboring non-metastatic cells have been found performing a k-NN search (implemented in fastNN) and then excluding pro-metastatic neighbors. The same procedure has been applied in the *‘Pooled-NN’* settings, with the only difference that PTs (and thus related pro-metastatic cells) have been pooled before NN search. Considering frequency differences between PT1 and PT5 pro-metastatic clones, we chose k=5 for PT1 *‘single-PT’* setting and k=10 otherwise.

### Gene modules analysis and signature scoring

Gene Modules (GMs) were found using the Hotspot [80], [81] method, fed with the raw gene expression matrix (only top 2000 HVGs) and the PCA space of all dataset cells. We used the ‘danb’ model. Hotspot GMs and curated gene signatures activation was scored for each cell using the AUCell method [81] with default parameters.

### Over-Representation and TF motifs enrichment analysis

Over Representation Analysis was performed with clusterProfiler [82], using as an input the top 50 genes from *‘some’* marker genes, DE and classification (see below) analyses, or the entire genes sets for Hotspot GMs. The top 50 enriched gene sets (ranked by qvalue) of each ORA were then considered for functional interpretation.

TF motifs enrichment analysis was performed with PASTAA (http://trap.molgen.mpg.de/PASTAA.htm) [27]. For a given gene list, this tool leverages promoter sequence and known TFs binding motifs information to identify the TFs that are putatively associated to those genes. We set the promoter range as an interval of 800 bp centered on the transcription starting site. We investigated only highly conserved sequences in promoters, and filtered only high-affinity TFs.

### Clonal distances and transcriptional heterogeneity analysis

Pairwise transcriptional dissimilarity of BC clones was quantified with various distance metrics.

Considering all clonal cell populations with size > 20 in some PT (n=119) as subsets of a metric space (i.e., the PCA space) we computed pairwise:

*i) centroid-centroid distances*. Considering two clones *X, Y* as disjoint subsets of the PCA space and *d* the Euclidean distance in this metric space, the centroid-centroid distance among *X, Y* was defined as:

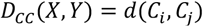

where each clone centroid is the arithmetic mean of its cell PCA coordinates;

*ii) Hausdorff distance*s [83]. Considering two clones *X, Y* as disjoint subsets of the PCA space and *d* the Euclidean distance in this metric space, the directed Hausdorff distance among *X, Y* was defined as:

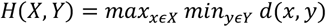

This asymmetric distance *H* was then symmetrized and normalized by the sum of the cardinalities of *X, Y*, ‖*X*‖ and ‖*Y*‖, yielding the (normalized) Hausdorff distance among the two clones:

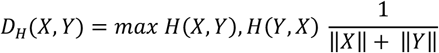

On the other hand, considering the same clonal cell populations as discrete, empirical probability distributions, we computed pairwise:

i. *Wassertein distances* [84] among clones’ cluster distributions. Here, each clone was represented by an 11 components vector, each storing the relative frequency of a the *i* −cluster in that clone;
ii. *Wassertein distances* among clones’ probability distributions over the PCA space. Each clone was represented by a matrix (with dimension: cells x PC_coordinates) and a vector (with dimension: cells) of uniform weights, each giving mass to the points of the PCA space occupied by the clone’s cells.

If X and Y are a pair of clones (in either of the above two representations), the Wassertein distance among *X, Y* was defined as:

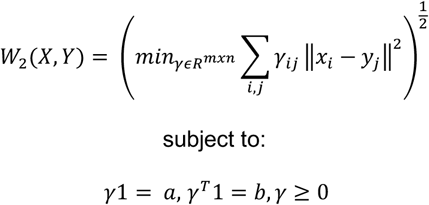

with γ the optimal transport plan transporting a unit mass from distribution *X* to distribution *Y*, and ‖*x*_*i*_ – *y*_*j*_‖^2^ the squared Euclidean distance among the *X, Y* clonal frequency components (case i)) or the PCA space coordinates of cells *x*_*i*_ and *y*_*j*_ (case ii)).

Transcriptional heterogeneity of the same clones was quantified with the SE of each cell population cluster distribution).

### scVI-based feature extraction

Raw HVGs expression values were fed to the variational autoencoder scVi [85]which was used an additional, denoised (sequencing-run, nUMIs and mitochondrial percentage were taken into account in the generative framework), non-linearly reduced (with n=30 features extracted) representation of the gene expression space. scVI hyperparameters were set as in [86]. scVI output was added to the other features considered for classification (i.e., PCA space coordinates and HVGs log-normalized expression data).

### Pro-metastatic/non-metastatic cells classification

Machine learning (ML) pipelines for (dominant) pro-metastatic vs other cells classification take full advantage of *numpy* [87], pandas [88], scikit learn [89] and XGboost [90] APIs. Pipelines from each classification scenario (i.e., separate or pooled PT1 and PT5, and across PTs) implement the following steps: *i) Data preprocessing*; *ii) Model selection*; and *iii) Performance evaluation* on held-out data.

#### i) Data preprocessing

Each classification pipeline started from a m x n features array ***X***, with m=number of cells of interests (e.g., PT1 cells), and n=dimension of the considered feature type, and a m x 1 binary vector ***y***, storing cell labels (i.e., ‘pro-metastatic’ or ‘other’ as derived from each cell clone membership). For each scenario and features choice (HVGs, PCA space or scVI cell embeddings), ***X*** and ***y*** arrays were split into training and test sets with a stratified shuffled split (80% of the total observation were assigned to training and 20% to test sets, respectively). Up-sampling of ‘pro-metastatic’ cells to a 1:10 total ratio was taken in consideration in some pipelines, but in most cases did not provide substantial benefit on final testing performances.

#### Ii) Model selection

XGBClassifier() and LogisticRegression() models were fit on training data as follows. First, key model hyper-parameters (see repo) were tuned by stratified 5-fold Cross Validation (CV) on training data, with the average F1-score over 5-fold CV as model performance metric (100 hyperparameters combinations were evaluated in each pipeline). Then, the optimal hyper-parameters combination was used to re-train the chosen model on the entire training set. Finally, the optimal model decision threshold was tuned considering again average 5-fold CV F1-scores.

#### iii) Performance evaluation on held-out data

The classification performance of a certain feature/model combination in a certain classification scenario was finally evaluated by the F1-score obtained on test set prediction.

*‘Single-PT’* and *‘Pooled-PT’* classification scenarios pipelines are all structured as described. On the contrary, *‘across_PT’* pipelines have a different training/test splitting logic, namely, an entire PT is used for training, while the remaining is used for testing.

Considered the achieved classification performances, we extracted lists of putatively relevant features only from *‘single-PT’* PT1 and PT5 best models (logistic regression fed with PCA space embeddings and logistic regression fed with HVGs expression data for PT1 and PT5, respectively). Normalized regression coefficients were used to rank features importance and prioritize putatively relevant genes: top 50 genes from PT5 best model were retained for further analysis and characterization, together with the top 5 genes with the highest loadings for each of the top 5 PCs ranked for PT1 best model.

### Survival analysis

ML pipelines for Survival Analysis take full advantage of *numpy*, pandas, scikit-learn, scikit-Survival [91] and lifelines [92] APIs. Pipelines from each survival task, implement the following steps: *i) Data preprocessing*; *ii) Enrichment of clinically relevant genes simulation*; *iii) Model selection;* and iv) *Multivariate analysis with known clinical factors*. Best signatures from ‘OS and all patients’ and ‘RFS and TNBC-only patients’ underwent an additional step v), namely *external cohort validation*. The following analytical workflow builds upon analysis from [93].

#### i) Data pre-processing

METABRIC z-scored expression values were downloaded from the cBioportal, together with clinical annotations. These data were wrangled to obtain a *samples x features* data matrix storing clinical (i.e., sample_id, patient_id, age, stage, subtype, OS and RFS survival time and status) and expression data for all diagnostic samples with no missing information for the included clinical annotations (n=1394). Stage IV and male patient samples were excluded from further analysis. A pool of putatively interesting genes (‘initial genes’) was assembled from the top 50 genes ranked within each DE and classification approaches (11 lists, 301 unique genes, 258 of which present in METABRIC microarray expression data).

#### ii) Enrichment of clinically relevant genes simulation

We initially assessed the clinical relevance of initial genes for each survival task by statistical simulation. We simulated 100 bootstrap samples from both initial genes and all-METABRIC genes (of note, results were independent of sample size, at least for tested values: 50, 75 and 100). Then, for all genes in these samples, we performed univariate survival analysis using the Cox Proportional Hazard Regression model (univariate Cox regression) implemented in CoxPHSurvivalAnalysis() from scikit-survival. For each sample we collected the number of genes scored as ‘clinically relevant’ (i.e., Harrel Concordance index (C-index) > 0.5 and Hazard Ratio (HR) > 1). Finally, we compared these simulated distributions with Wilcoxon’s Test.

#### >iii) Model selection

The best molecular predictor for each survival task (i.e., gene signatures) was learned from the pre-processed METABRIC dataset subsetted only for initial genes and task-associated samples (e.g., only TNBC samples for TNBC OS and RFS survival tasks). First, univariate Cox regression was applied to filter only clinically relevant genes. This step yielded one filtered gene set for each task **(Suppl. Tables 6)**. These sets were then used to feed multivariate Cox regression models with forward feature selection. Specifically, for each task and related filtered gene set, a ‘seed’ gene was chosen (i.e., the gene showing highest rankings for both CI and HR in the univariate setting). Then, all the other genes were added to the initial seed, and multivariate Cox regression was used to test the predictive performance of all these 2-gene signatures. The best (highest C-index) 2-gene signature was then chosen to undergo another iteration, and so on. The updating scheme stopped at CI convergence.

For each survival task, this simple modeling framework achieved equal or better performance than all the other tested approaches, namely elastic nets Cox regression and gradient boosted models implemented in scikit-survival. This pattern was invariably observed starting from both initial and filtered gene sets. Performance of all modeling approaches was evaluated with 5-fold CV.

#### iv) Multivariate analysis

We tested the independence of each signature predictive power from other known clinical covariates with a multivariate analysis on METABRIC data. Specifically, for each signature we calculated the following score:

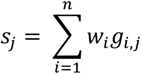

where for signature *sig* := {*g*_*i*_}, *s*_*j*_ is the jth-sample score; *g*_*i,j*_ is the expression value of the signature gene *g*_*i*_; *w*_*i*_ is *g*_*i*_ regression coefficient in the multivariate model that generated *sig* and *i* = 1, …, *n* genes in *sig*. For each signature multivariate Cox PH regression analysis was then performed with the CoxPHfitter() function implemented in lifelines, including (scaled) signature score, (scaled) age, stage and optionally (if not a TNBC-only task) one-hot-encoded subtype as model covariates. Additionally, Kaplan-Meier (KM) Curves were made with the kaplan_meier_estimator() function from sci-kit survival, assigning patient into ‘high’ and ‘low’ signature expression categories if their signature score was above or below the median value, respectively. The Log-rank sum test was used to quantify each signature score stratification capacity.

#### iii) External cohort validation

External cohort datasets were downloaded from GEO (see Fig. S5 caption for GEO accession numbers) and preprocessed as for the METABRIC dataset. PCA performed on concatenated datasets (Z-scored initial genes expression values) were used to visually identified potential dataset specific sources of variation, potentially preventing merging (data not shown). KM curves were made as previously described.

### Target validation in vitro

The shRNAs (sequences are reported in **Suppl. Table 9**) were cloned in the pRSI17-U6-sh-UbiC-TagGFP-2A-Puro Plasmid as previously described. Two distinct shRNA sequences were designed per each target. A scrambled shRNA (Cellecta) [94] was used as a control.

### Quantification of gene silencing

To quantify the extent of gene silencing, MDA-MB-231 cells were infected with either the scrambled or the targeting shRNA and selected by puromycin (2.5μg/ml for 72h; Vinci-Biochem). The extent of gene expression downregulation was quantified 7 days after infection by qPCR (7500 Real-Time PCR System), using the scrambled shRNA as calibrator. The list of primer is provided in **Suppl. Table 9**.

### Cell proliferation and cell cycle assay

To assess the effect of the silencing on cell proliferation, 300,000 MDA-MB-231 were plated 7 days after infection in three independent plates. Cells were counted respectively after 72, 168, and 240h after plating. The computed number of cells was normalized on the initial number of plated cells. Cells were split once they reached ∼80% confluence and the splitting factor was considered throughout the experiment. To assess the effect of gene silencing on cell cycle distribution, MDA-MB-231 were harvested 240h after plating, fixed with 70% cold ethanol (Panreac Applichem) in PBS (Thermo Fisher Scientific), and stored at 4°C. After 8h, cells were centrifuged and resuspended in PBS (Thermo Fisher Scientific). DAPI (Sigma-Aldrich) was then added at the final concentration of 50µM overnight at 4°C. Cell cycle distribution was investigated by FACS analysis.

### Cell migration assay

To assess the effect of gene silencing on cell migration, MDA-MB-231 underwent wound healing assay 7 days after infection. For this aim, 200,000 cells/well were plated in a 12-well plate. The day after plating, the cell monolayer was scratched with a 20μl sterile pipette tip and washed with DMEM (Euroclone). Then, a photo of the wound was taken by EVOS® FL Cell Imaging System (Thermo Fisher Scientific) and the photographed region was marked. Cells were cultured in DMEM (Euroclone) for additional 8h so to rule out the role of proliferation in this assay. Ultimately, a second photo of the region was taken, and the migration extent was computed as wound closure (i.e., the difference in terms of area devoid of cells between the first and the second time point). The migration in the sample infected with the targeting shRNA was normalized on the migration in the sample infected with the scrambled shRNA.

### Target validation in mouse

MDA-MB-231 were initially infected with H2B-GFP reporter construct to allow their identification *in vivo*. Upon infection, cells expressing high levels of GFP were isolated from dissociated tissues via FACS using a BD FACSAria II. Sorted cells were subsequently infected with either shRNA targeting the gene of interest or scrambled shRNA and selected by puromycin (2.5μg/ml for 72h; Vinci-Biochem). Double-infected cells were cultured in vitro for 4 additional days to allow a stable gene silencing. Prior to transplantation, cells were further infected with firefly luciferase pLenti CMV Puro LUC (w168-1) reporter vector at MOI = 5, so to ensure that most cells expressed luciferase. Per each mouse, 200,000 MDA-MB-231 cells were resuspended in 30 μl of PBS (Thermo Fisher Scientific) and growth-factor-reduced Matrigel (Corning) in a ratio of 1:1. Cells were injected by intra-ductal transplantation in the ninth mammary gland of 8–12-week-old female NSG mice without prior clearing of the fat pad. MDA-MB-231 cells expressing a scrambled shRNA were transplanted into 7 mice, while MDA-MB-231 expressing an shRNA against the gene of interest were transplanted into 14 mice (7 per each different gene-targeting construct). Tumor growth was weekly monitored by IVIS-Lumina *in vivo* imaging system (Perkin Elmer) and tumor volume was estimated as previously described [63]. Tumors were resected ∼40 days post injection (once they reached a volume of 0.5-1 cm^3^), and mice were sacrificed 21 days after PT resection. Lungs were harvested and 200μm thick slices were generated by Leica VT1200 semiautomatic vibrating microtome (Leica). The presence of H2B-GFP positive metastatic cells was investigated by Eclipse Ti2 inverted microscope (Nikon). In line with previous works [95], we applied to each image a custom macro aimed to determine the area occupied by metastatic cells in each lung section. In this macro, the perimeter of the section was traced manually, and the area of the slice calculated consequently. The image was filtered and thresholded to remove all background and create a binary image. Then, the total area occupied by metastatic cells was computed. The obtained value was divided by the total area of lung slice, and this value was further divided by the volume of the PT measured before resection [95] to obtain the metastatic burden. Ultimately, the metastatic burden was normalized to the value in the scrambled shRNA cohort and the corresponding values in the targeting shRNA cohorts were computed consequently.

### Target validation in zebrafish

MDA-MB-231 were initially infected with H2B-GFP reporter construct to allow for their identification *in vivo*. FACS-Sorted H2B-GFP expressing cells from dissociated tissues were subsequently infected with either shRNA targeting the gene of interest or scrambled shRNA and selected by puromycin (2.5μg/ml for 72h; Vinci-Biochem). Double-infected cells were cultured *in vitro* for 4 additional days to allow a stable knock-down of the genes of interest. Then, cells were resuspended in PBS (Thermo Fisher Scientific) to the final concentration of 100,000cells/μl. Prior to injection, 2 dpf larvae were manually dechorionated and anesthetized with 0.016% tricaine (Ethyl-3-aminobenzoate methane-sulfonate salt, MS-222, Sigma-Aldrich) in E3 water medium supplemented with PTU. Transplantation was performed under Olympus SZX9 stereomicroscope (Nikon) using the pneumatic Picospritzer III micro-injector (Parker Instrumentation). Customized micro-injection needles were prepared pulling GC100T-15 borosilicate glass capillaries (1,0 mm outer diameter/0,78 mm inner diameter; Harvard Apparatus) using a P-97 micropipette puller device (Sutter Instruments) and the following customized parameters: heat=505; pull=60; velocity=60; time=100; air pressure=300. Injection capillary was loaded with 10 µl of cell suspension and installed on a manual micromanipulator (Narishige) in order to ensure fine movements. The tip of the needle was manually broken off with fine tweezers in order to obtain a tip opening diameter of 5-10 µm. Anesthetized larvae (∼100 animals per cohort) were injected into the PVS with 5nl of cell suspension, corresponding to ∼500 cells. Following transplantation, larvae were transferred to E3 water + PTU, left to recover at 28.5°C for 30 minutes, and then selected for correct transplantation with a Nikon SMZ25 stereomicroscope. Dead, abnormal, non-injected or mis-injected larvae were discarded. Correctly injected xenografts were transferred to a 34°C incubator for the remaining experimental time-window, to meet the optimal temperature requirements for fish and mammalian cells. At 1 and 4 dpi fish were subjected to a second screening to discard dead and abnormal animals. Remaining animals were maintained in 48-multi-well plates for subsequent analyses. The incidence of metastasis at both time points was calculated as the fraction of larvae displaying at least one (>5 cells) metastasis over the total number of larvae that survived till that time point. To quantify PT cellularity, 4 dpi transplanted animals were anesthetized with 0.016% tricaine in E3 water + PTU, included in 0.8% low-melting point agarose dissolved in E3, and placed under a Leica SP8 AOBS confocal microscope. Z-stacks tile scan images of the whole larva were acquired with a 10x/0.3 N.A. dry objective. Magnifications of the anterior portion of the larva were acquired with a 25x/0.95 N.A. water immersion objective, with 5 µm z-stack interval, in resonant scanner modality. Samples were simultaneously excited with 488 nm and 561 nm lasers to acquire signals from GFP^+^ cancer cells and DsRed^+^ zebrafish endothelial cells. Quantification of PT cellularity was performed with Arivis Vision 4D 3.5.0 software, using a customized pipeline. Images were first denoised via “particle enhancement” and “gaussian filter” functions and then segmented via “blob finder” option on the GFP channel. All the segmented objects that met given requirements of volume (volume > 100 µm^3^) and sphericity (sphericity between 0.4 and 1) were considered as H2B-GFP^+^ nuclei.

### Statistics

All statistical comparisons were evaluated with the Wilcoxon-Mann-Whitney test, with no multiple hypothesis testing correction of p-values unless otherwise stated (see figure legends). All boxplots present the quartiles of the distribution and whiskers show the rest of the distribution.

---

**Fig. S1.**
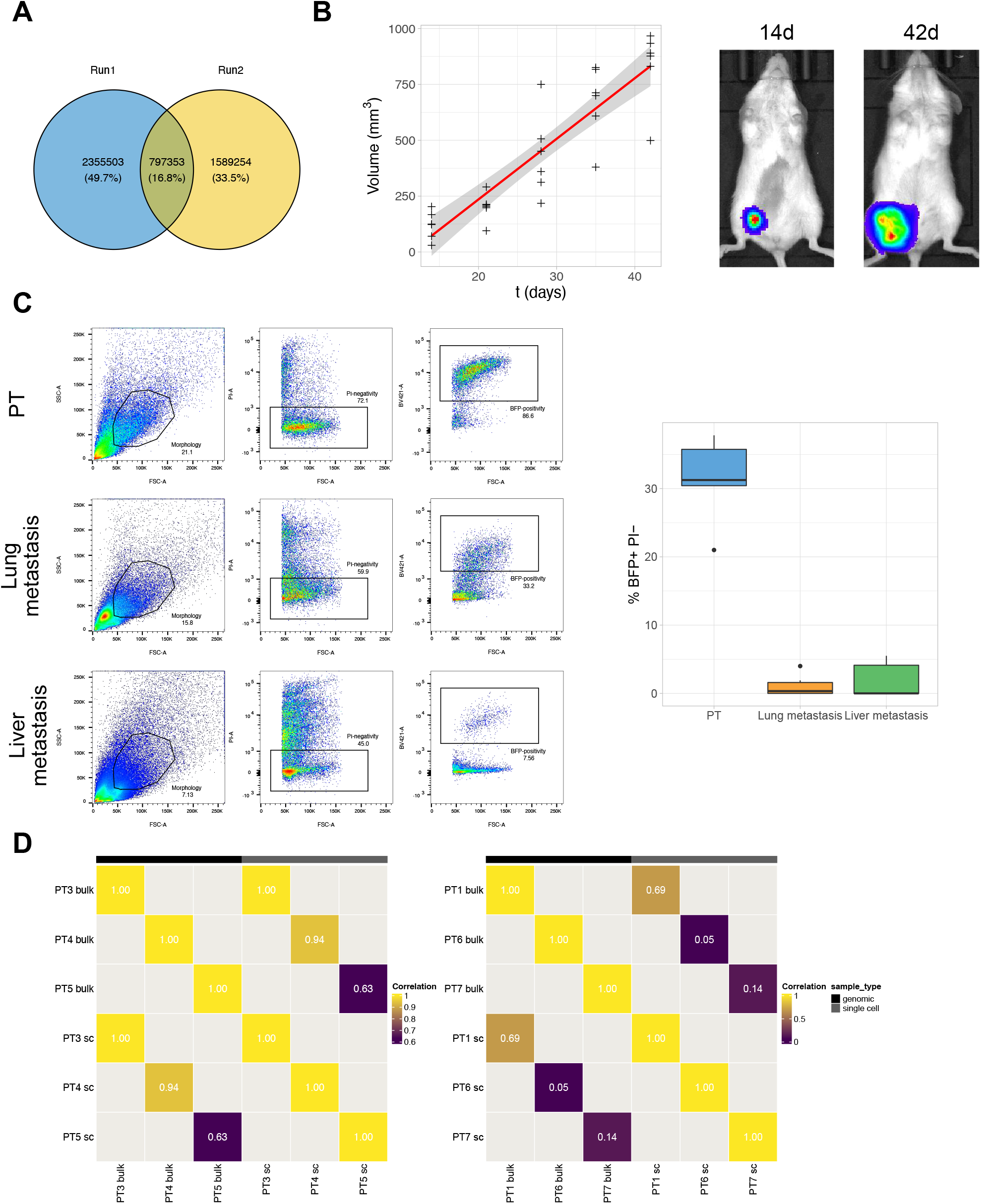
Pro-metastatic clones are rare and under-represented in primary tumors, and form monoclonal metastases. (A) Numbers of total and overlapping reads of two sequencing-runs of the Perturb-seq GBC library. (B) *In vivo* progression of luciferase-expressing MDA-MB-231 PTs measured by IVIS at the indicated time points. Kinetics of growth in 6 distinct mice (left panel) and representative images of one mouse at two different time points, as indicated (right panel). The blue curve represents the fitted trend (linear regression) of tumor volume over time, while the grey area corresponds to the confidence interval (C) FACS-sorting of alive (Propidium iodide (PI)-negativity) and GBC-barcoded (BFP-positivity) MDA-MB-231 cells from PTs, lung metastases and liver metastases, as indicated. Gating strategies for the isolation of tumor cells (left panel): cells were gated sequentially by complexity (forward *vs* side scatter), PI-negativity and BFP-positivity, as indicated (numbers indicate percentages of cells in each gate). Average percentage of BFP+ PI-cells recovered from each sample (right panel). (D) Pearson pairwise correlation of clone frequencies of the indicated genomic (bulk) and scRNA-seq (Sc) samples. Numbers within the squares indicate the pairwise Pearson correlation coefficient.

**Fig. S2.**
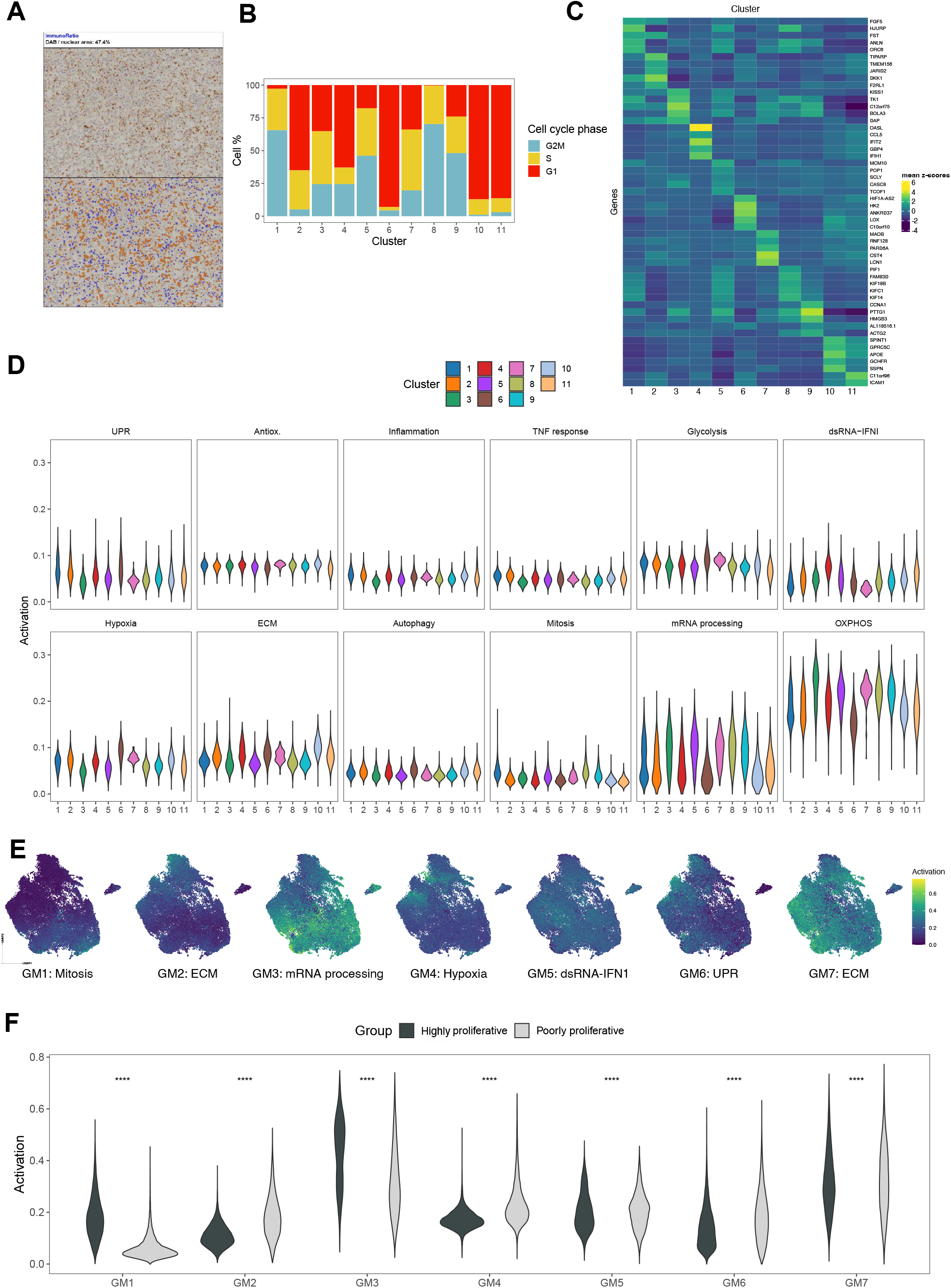
Dominant pro-metastatic clones are characterized by the same intra-clonal heterogeneity of all other PT clones. (A) Immuno-histochemical analyses of Ki67 expression. Sections from paraffin-embedded PT were stained with anti-Ki67 antibody and hematoxylin. Ki67-positive cells were revealed by brown diaminobenzidine (DAB) staining. Images were processed with the ImmunoRatio ImageJ plug-in. The upper panel shows one representative DAB-hematoxylin staining field, the lower is the corresponding pseudo-colored image showing the segmented staining components (i.e., Ki67-positive nuclei in brown and Ki67-negative nuclei in violet). The fraction of Ki67+ cells was quantified through calculation of DAB-stained area/total nuclear area. (B) Cell-cycle profile of the 11 clusters. Percentages of G1, S and G2M cells are indicated by color code. (C) Marker gene analysis of the 11 clusters. Per each cluster, the top 5 marker genes are shown, when available. Color code represents the level of expression of each gene, as indicated. (D) Violin plots showing expression of gene signatures associated to mitosis (GOBP_MITOTIC_NUCLEAR_DIVISION), mRNA processing (GOCC_SPLICEOSOMAL_TRI_SNRNP_ COMPLEX), OXPHOS (GOBP_OXIDATIVE_PHOSPHORYLATION), UPR (GOBP_CELLULAR_RESPONSE_TO_UNFOLDED_PROTEIN), hypoxia (GOBP_CELLULAR_RESPONSE_TO_ OXYGEN_LEVELS), autophagy (GOCC_AUTOPHAGOSOME), dsRNA-IFN1 signaling (GOBP_CELLULAR_RESPONSE_TO_DSRNA), TNF response (GOBP_RESPONSE_TO_TUMOR_NECROSIS_FACTOR), ECM remodeling/ interaction (GOCC_COLLAGEN_CONTAINING_EXTRACELLULAR_MATRIX), glycolysis (HALLMARK_GLYCOLYSIS), oxidative stress response (GOBP_RESPONSE_TO_OXIDATIVE_STRESS), and inflammation (GOBP_RESPONSE_TO_MOLECULE_OF_BACTERIAL_ORIGIN) across clusters. (E) UMAP representation of the expression of the 7 GMs across the 37,801 single cells, with pathways associated to each GM reported below each UMAP. Pathways were identified by the top 1-2 gene sets in the over-representation analysis performed on the genes of each GM. (F) Violin plot showing expression of genes associated to each GM in all Group 1 (1, 5, 8, and 9) *vs* all Group 2 (2, 3, 4, 6, 7, 10 and 11) clusters (Wilcoxon-Mann-Whitney test; **** p < 0.0001).

**Fig. S3.**
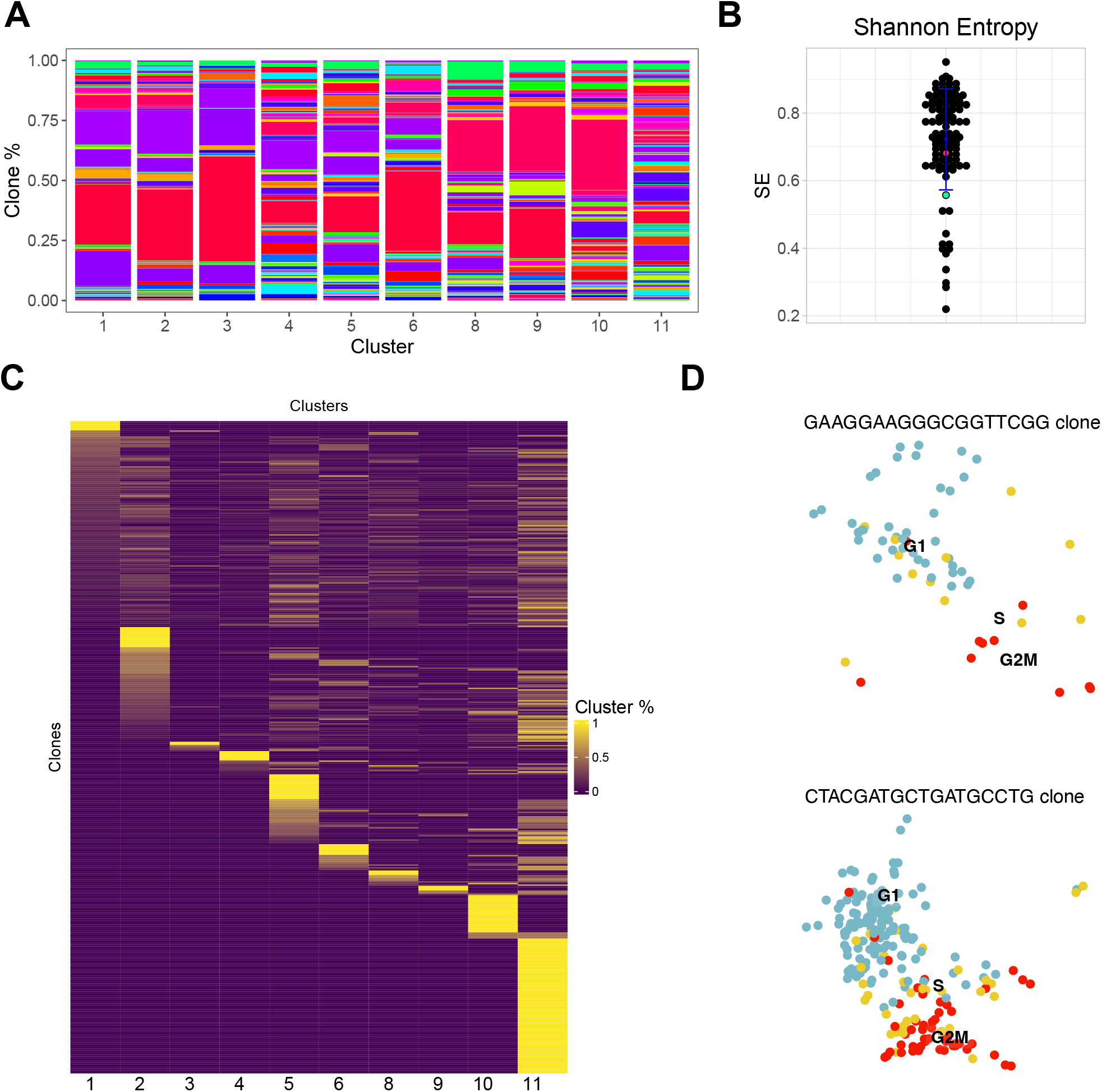
Dominant pro-metastatic clones are characterized by the same intra-clonal heterogeneity of all other PT clones. (A) Clonal composition of clusters. Only clones from PTs are reported. Each color identifies a specific clone as in Fig. 1 and 3. (B) Cluster distribution Shannon Entropy (SE) of each clone (mean and standard deviation in blue; PT1 and PT5 dominant pro-metastatic clones in green and red). (C) Clonal distribution across clusters. Only clones from PTs are reported (clones >20 cells). Each row identifies a clone, and the color code describes the fraction of each clone (defined as Cluster %) contained in each cluster (column). (D) UMAP representation showing the cell cycle phase distribution of pro-metastatic clones in PT1 (left panel) and PT5 (right panel).

**Fig. S4.**
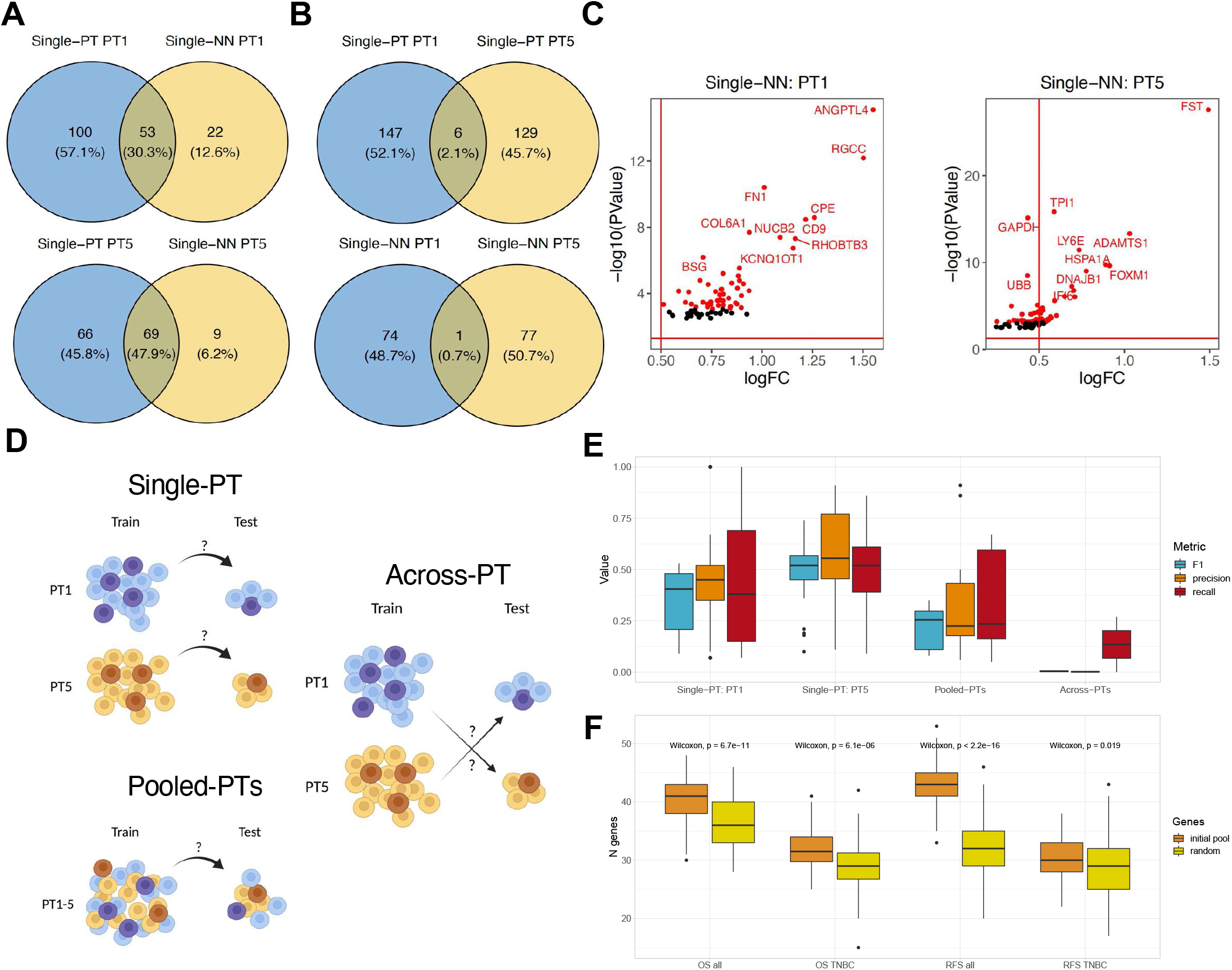
Dominant pro-metastatic clones are distinguished by up-regulation of genes that predict disease progression in BC patients. (A) Venn diagram representing the top 50 genes in the ‘single-NN’ and ‘single-PT’ settings (upper panel PT1, lower panel, PT5). (B) Venn diagram indicating the number of shared and unique DEGs identified in the ‘single-NN’ and ‘single-PT’ settings between PT1 and PT5 pro-metastatic clones. The top 50 DEGs (ranked by FDR) are colored in red. (C) Volcano plot representing PT1 and PT5 pro-metastatic DEGs identified in ‘single-NN’ setting. Per each DEG, we reported the corresponding logFC (x-axis) and -log10(Pvalue) (y-axis). The top 50 DEGs (ranked by FDR) are colored in red. (D) Schematic representation of the classification approach. Left panels: the model was trained in three different different contexts (single PT1, single PT5, or pooled PT1-PT5; train) and tested (test) on the same samples, as indicated. Right panels: training was performed on PT1 and tested on PT5, and viceversa. (E) Box-plot of precision, recall, and F1 score value in single PTs (PT1 or PT5), pooled-PTs (PT1-PT5) and across-PT predictions. (F) Box-plot comparing numbers of genes with HR>1 for the four survival tasks in our initial gene list (100 random samples of 100 genes from the list of 301 genes) and in all the genes identified in the METABRIC study (100 random samples of 100 genes from the list of all the genes identified in the METABRIC study; Wilcoxon-Mann-Whitney test, p-values are reported).

**Fig. S5.**
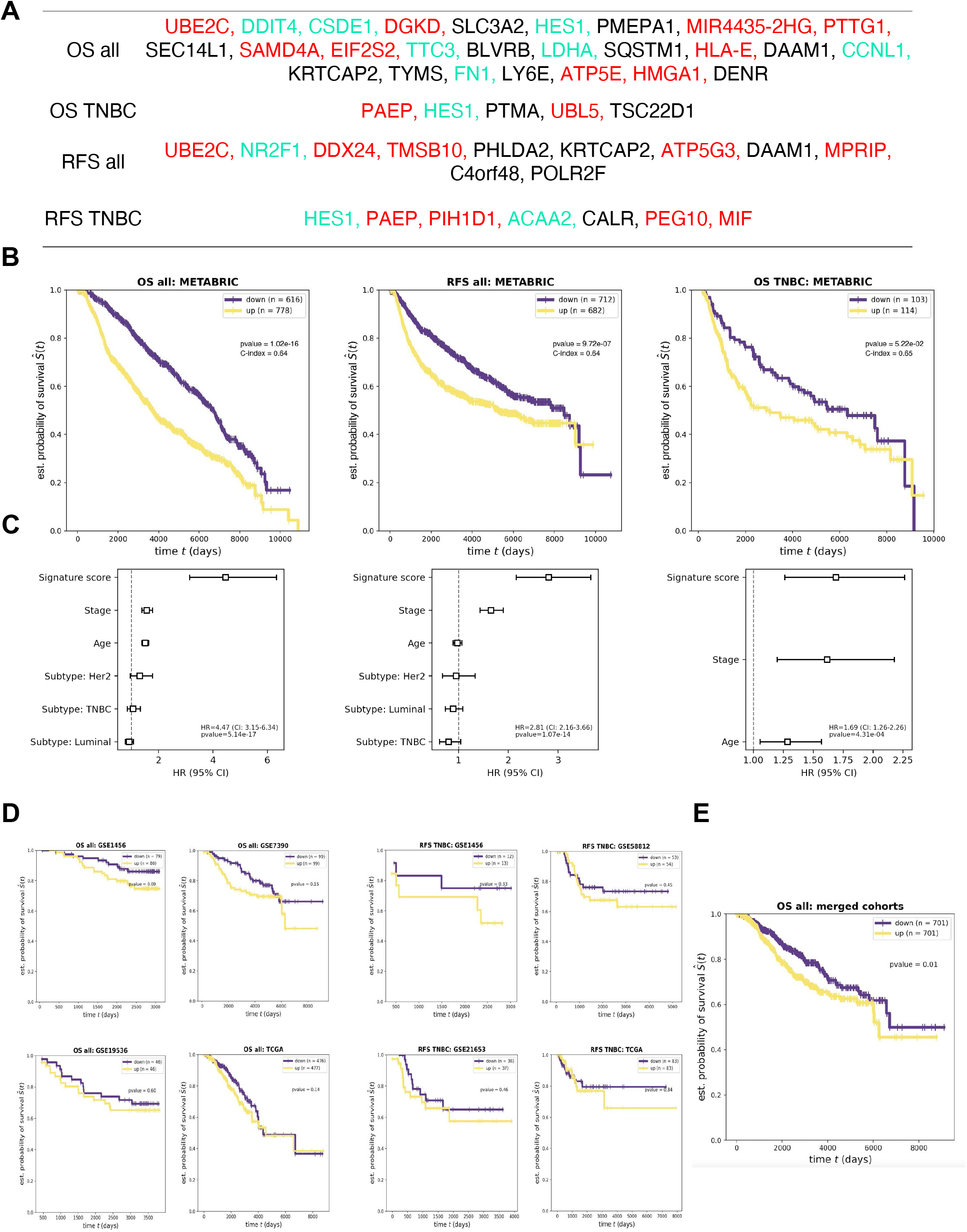
Dominant pro-metastatic clones are distinguished by up-regulation of genes that predict disease progression in BC patients. (A) List of genes composing the 4 gene signatures. PT1 and PT5 pro-metastatic genes are respectively reported in green and red. Pro-metastatic genes expressed by both clones are reported in black. (B) Kaplan-Meier curve of respectively OS survival in all BC patients, RFS in all BC patients, and OS survival in TNBC patients according to high or low expression of the corresponding signature in the training set (METABRIC; CI and p-values calculated by two-sided log-rank test are reported). The y-axis reports the estimated probability of survival (Ŝ) over time (i.e., the number of subjects surviving divided by the number of patients at risk), while the x-axis reports the time (in days). (C) Multivariate analysis of respectively OS all, RFS all, OS TNBC gene signature independence of other confounding factors. We computed the HR of signature (sig_score, see M&M), tumor stage, age at diagnosis, and (optionally) subtype separately and using the other factors as model covariates. (D) Kaplan-Meier curve of OS in all BC patients according to high or low expression of the corresponding signature in 4 distinct clinical studies (GEO accession number are reported above the corresponding curve). p-values were calculated by a two-sided log-rank test. (E) Kaplan-Meier curve of RFS in TNBC patients according to high or low expression of the corresponding signature in 4 distinct clinical studies (GEO accession number are reported above the corresponding curve). p-values were calculated by a two-sided log-rank test. (F) Kaplan-Meier curve of OS in all BC patients according to high or low expression of the corresponding signature in the test set (Multiple merged cohorts, the GEO accession numbers of merged clinical studies are reported in Fig. S5D; p-values calculated by a two-sided log-rank test is reported).

**Fig. S6.**
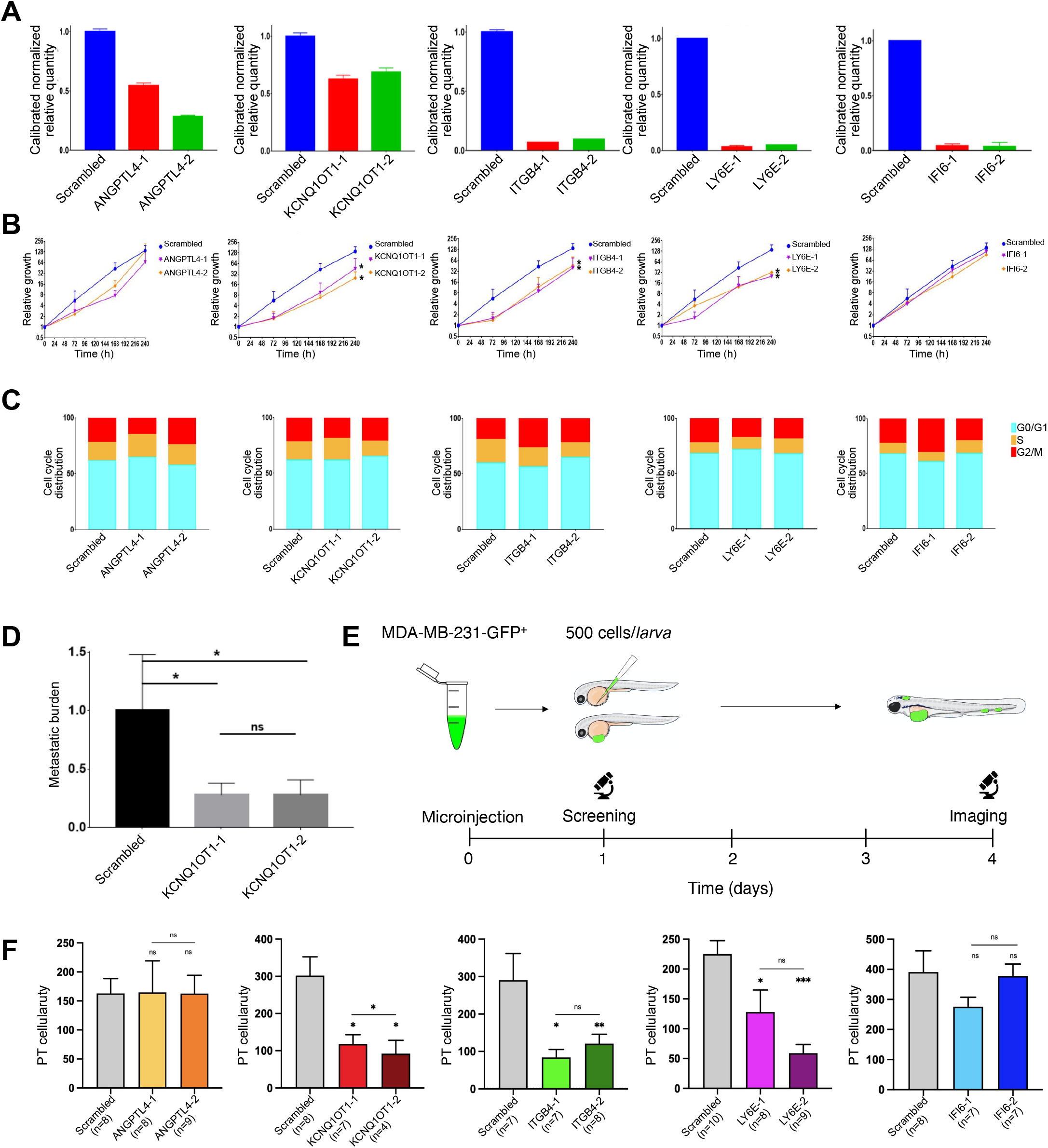
Silencing of single pro-metastatic genes dramatically reduces metastatization *in vivo*. **(A)** Extent of gene silencing, measured by qPCR, upon MDA-MB-231 infection with either the scrambled or the two targeting shRNAs per each validated gene. **(B)** Analysis of the effects of gene silencing on *in vitro* proliferation for up to 10 days (n = 3 per each panel; two-tailed T-test, *p < 0.05; ** p < 0.01). Cells were manually counted at 72, 168, and 240 hours after initial palting. Only alive cells (discriminated by erythrosin B-negativity) were counted. The computed number of cells was normalized on the initial number of plated cells. Cells were split once they reached ∼80% confluence and the splitting factor was considered throughout the experiment. **(C)** Analysis of the effects of gene silencing on cell cycle distribution 10 days after *in vitro* plating. Cells were stained with DAPI and their distribution across cell cycle phases was estimated by FACS analysis. **(D)** Quantification of metastatic burden (see M&M) in mouse carrying MDA-MB-231 cells infected with either the scrambled or the two KCNQ1OT1-shRNAs (Scrambled cohort n = 7; KCNQ1OT1_1 cohort n = 7; KCNQ1OT1_2 cohort n = 7; two-tailed T-test, ** p < 0.01). **(E)** Schematic representation of *in vivo* validation experiment in zebrafish. ∼500 MDA-MB-231 constitutively expressing the H2B-GFP and the shRNA targeting the gene of interest were injected in the PVS of 2 days post fertilization zebrafish larvae. Metastatic progression was monitored at 4 dpi by fluorescence microscopy: we considered as metastases uniquely the nodules > 5cells which colonized the caudal portion of zebrafish larvae. **(F)** Bar plots of PT cellularity (see M&M) in zebrafish larvae computed 4 days after transplantation (mean + SEM are displayed, Kruskal-Wallis test, *p < 0.05; ** p < 0.01; *** p < 0.001). Images of the apical zebrafish portion were segmented to identify H2B-GFP nuclei in the PT and the total number of nuclei in a zebrafish z-stack were computed to define PT cellularity.

